# A global map of wood density

**DOI:** 10.1101/2025.08.25.671920

**Authors:** Fabian Jörg Fischer, Jérôme Chave, Amy Zanne, Tommaso Jucker, Alex Fajardo, Adeline Fayolle, Renato Augusto Ferreira de Lima, Ghislain Vieilledent, Hans Beeckman, Wannes Hubau, Tom De Mil, Daniel Wallenus, Ana María Aldana, Esteban Alvarez-Dávila, Luciana F. Alves, Deborah M. G. Apgaua, Fátima Arcanjo, Jean-François Bastin, Andrii Bilous, Philippe Birnbaum, Volodymyr Blyshchyk, Joli Borah, Vanessa Boukili, J. Julio Camarero, Luisa Casas, Roberto Cazzolla Gatti, Jeffrey Q. Chambers, Ezequiel Chimbioputo Fabiano, Brendan Choat, Edgar Cifuentes, Georgina Conti, David Coomes, Will Cornwell, Javid Ahmad Dar, Ashesh Kumar Das, Magnus Dobler, Dao Dougabka, David P. Edwards, Urs Eggli, Robert Evans, Daniel Falster, Philip Fearnside, Olivier Flores, Nikolaos Fyllas, Jean Gérard, Rosa C. Goodman, Daniel Guibal, L. Francisco Henao-Diaz, Vincent Hervé, Peter Hietz, Jürgen Homeier, Thomas Ibanez, Jugo Ilic, Steven Jansen, Rinku Moni Kalita, Tanaka Kenzo, Liana Kindermann, Subashree Kothandaraman, Martyna Kotowska, Yasuhiro Kubota, Patrick Langbour, James Lawson, André Luiz Alves de Lima, Roman Mathias Link, Anja Linstädter, Rosana López, Cate Macinnis-Ng, Luiz Fernando S. Magnago, Adam R. Martin, Ashley M. Matheny, James K. McCarthy, Regis B. Miller, Arun Jyoti Nath, Bruce Walker Nelson, Marco Njana, Euler Melo Nogueira, Alexandre Oliveira, Rafael Oliveira, Mark Olson, Yusuke Onoda, Keryn Paul, Daniel Piotto, Phil Radtke, Onja Razafindratsima, Tahiana Ramananantoandro, Jennifer Read, Sarah Richardson, Enrique G. de la Riva, Oris Rodríguez-Reyes, Samir G. Rolim, Victor Rolo, Julieta A. Rosell, Sassan Saatchi, Roberto Salguero-Gómez, Nadia S. Santini, Bernhard Schuldt, Luitgard Schwendenmann, Arne Sellin, Timothy Staples, Pablo R Stevenson, Somaiah Sundarapandian, Masha T van der Sande, Hans ter Steege, Shengli Tao, Bernard Thibaut, David Yue Phin Tng, José Marcelo Domingues Torezan, Boris Villanueva, Aaron Weiskittel, Jessie Wells, S. Joseph Wright, Kasia Zieminska, Alexander Zizka

## Abstract

Wood density influences how quickly woody plants grow, how long they live and how much carbon they store, yet its global variation remains poorly mapped. Here we combined 109,626 wood density measurements from 16,829 species with 300,949 vegetation plots to produce a km-scale map of community-weighted wood density for every woody biome. Our model led to a prediction accuracy 32–51 % higher than previous global products, and a 1.8–3.7-fold wider wood density range (0.28–1.00 g cm^−3^; global mean: 0.57 g cm^−3^) than previously assumed. Spatial cross-validation showed low bias (±2.5 % of the mean), and uncertainties decreased from 20% in poorly sampled drylands and boreal regions to 5% in data-rich temperate forests. Mean annual temperature was the best predictor of community-weighted mean wood density, increasing by 0.01 g cm^−3^ for every 1°C change. We deliver a low-bias, high-resolution wood density layer for Earth system models, together with spatially explicit error maps. This study represents a major step forward for carbon accounting and trait-based forecasts of vegetation change.

---

The world’s woody vegetation plays a major role in the global carbon cycle. It sequesters and stores carbon and, in doing so, slows down anthropogenic climate change^1^. Monitoring and protecting carbon stocks is an essential mitigation strategy^2,3^ and has been enshrined in policy commitments^4^. However, we still lack a reliable global quantification system for carbon storage in live vegetation^5^. Such a system requires wall-to-wall quantification of the volume of woody plants and the density of their wood (dry mass per unit green volume; g cm^-3^). Volume estimates are rapidly improving thanks to a growing array of air- and spaceborne sensors^6–8^, but measuring wood density relies on species identification and tissue extraction on the ground^9^. This process is labor-intensive and leaves massive data gaps: species composition and mean wood densities are mostly unobserved by satellites, potentially biasing carbon estimates^10^. It is thus critical to bridge gaps in wood density information with robust statistical models.

Several wood density maps have been produced at global^11–13^ and regional^14–16^ scales, but three types of inconsistencies are hampering efforts towards comparability across maps. First, wood density cannot be measured for every single plant in a focal area, meaning values must be estimated^17^. Studies differ in how they generate estimates, what databases they draw on and how they standardize wood density values^18^. Second, studies use different definitions and datasets to aggregate wood density at the level of ecological communities. To account for species that are more abundant or biomass-rich than others, community means are usually obtained by weighting the contribution of all individuals in a focal area equally or in proportion to their cross-sectional (or basal) trunk area^19^. These community-weighted means appear robust to the choice of weights^15^, but require comprehensive field inventories, which are rare, so studies often use geographically imbalanced inputs^11^, include unevenly and partially sampled communities^12,13^, or average directly across species occurrence records^16^. Third, modelling wood density across spatial scales represents a great challenge. At broad scales, community-weighted mean wood density varies with floristic composition^19^, reflecting shifts in ecological strategies between taxonomic groups (e.g., angiosperms vs. gymnosperms), life forms (e.g., succulents vs. non-succulents) and among species within these groups. All else being equal, plants with high wood density are expected to be shorter and grow slower in volume than less dense plants^20–22^. During early ecological succession, these high wood density plants are at a disadvantage in the competition for light, but in the long run, high wood density reduces hydraulic and mechanical risks and increases competitive ability^23^. We thus expect low mean wood densities in fast-turnover environments, such as under frequent cycles of disturbance or recovery, or on fertile soils^24–27^, and high wood densities in hot and dry areas^28–30^. However, it is a major challenge to select appropriate predictors and account for spatial autocorrelation in model calibration and validation^31^.

Here, we map community-weighted mean wood density at 1 km^2^ resolution globally, mindful of the three challenges outlined above. To produce the map, we used a major update of the Global Wood Density Database (GWDD v.2)^32^, which contains over 6x times as many wood density records (109,626) as the original and over 2x as many species (16,829), all rigorously quality-checked and standardized^33^. We then compiled a geographically balanced dataset of 300,949 vegetation inventories with detailed floristic information across all woody biomes and used it to compute community-weighted wood density, here defined as mean wood density weighted by basal area. Third, we mapped community-weighted mean wood density with random forest models across the globe at 1 km^2^ resolution. We selected 10 environmental predictors with biologically meaningful links to wood density variation, thoroughly validated the wood density map, and compared it with three published global maps^11–13^ as well as a regional map^15^. Our results represent a major step forward in creating a reliable global carbon quantification system and improving climate change assessments.

## Results

### Global distribution of community-weighted mean wood density

Community-weighted mean wood density varied more than 3-fold from 0.28 g cm^-3^ to 1.00 g cm^-3^ across the globe, with a mean of 0.57 g cm^-3^ and standard deviation of 0.104 g cm^-3^ (95% confidence interval, 95%CI: 0.40 to 0.77 g cm^-3^, n_pixel_ = 86.7 million, Fig. 1a). Values displayed a bimodal distribution with one mode at 0.42 g cm^-3^, reflecting gymnosperm-dominated forests at high latitudes, and a second mode at 0.64 g cm^-3^, reflecting tropical forests and savannas (Fig. 1a, Table S1). Dry biomes, such as deserts and xeric shrublands had the highest average densities at 0.66 g cm^-3^ (95%CI: 0.47 to 0.79 g cm^-3^, Table S1), but also large biogeographic variation. Our map predicted much higher average wood densities of 0.76 g cm^-3^ (95%CI: 0.64 to 0.87 g cm^-3^) than the dry-biome average across xeric environments in Australia and much lower densities of 0.46 g cm^-3^ (95%CI: 0.40 to 0.63 g cm^-3^) in succulent-dominated Baja California. In the boreal zone, larch-dominated forests of Northeast Siberia exceeded the boreal average by ca. 20%, with wood densities of 0.50 g cm^-3^ (95%CI: 0.43 to 0.63 g cm^-3^).

**Fig. 1:**
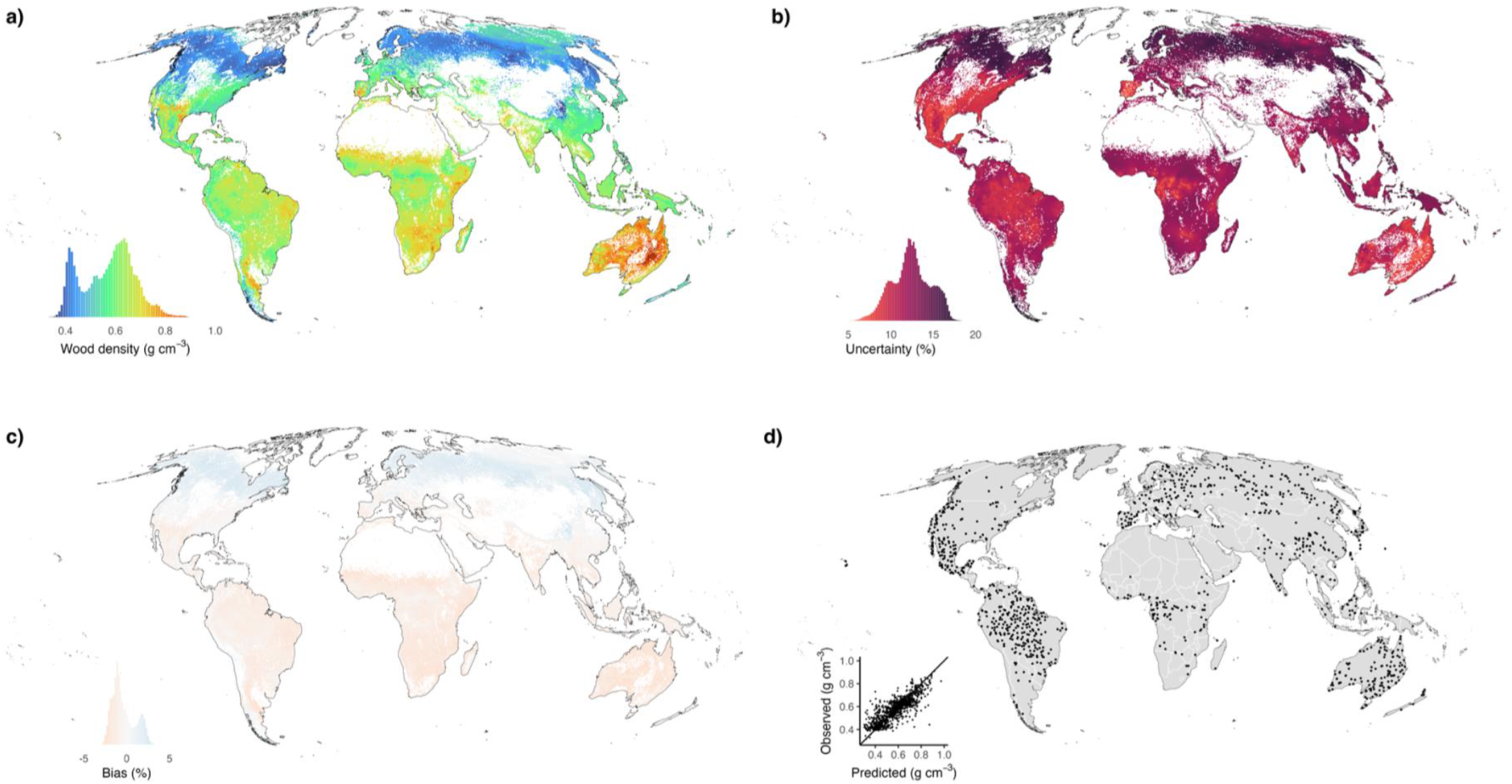
Global maps of wood density, prediction errors and validation plots. Shown is the predicted global distribution of community-weighted mean wood density (g cm^-3^, a), relative prediction uncertainty (% of predicted wood density, b), relative prediction bias (% of predicted wood density, c) and the validation plots (d). Maps have a 1 km resolution in Mollweide equal area projection. In panels a-c, insets show both the colour scale and the histogram of values. In panel d, the inset shows predicted vs. observed wood densities for all validation plots in basic cross-validation (i.e., not accounting for distance to training data). Areas without tree or shrub cover are masked. For absolute uncertainty and bias maps see Fig. S5. In panels b and c, colour scale breaks have been matched to cover the same 5% intervals.

### Map validation and error assessment

To create spatially explicit estimates of uncertainty and bias, we performed a spatially stratified cross-validation. We selected a geographically balanced subset of 1,000 plots (median area: 0.64 ha, Fig. 1d), compared modelled and observed wood densities, and used the results to map uncertainty (Fig. 1b) and mean error (or bias, Fig. 1c). Overall bias was low, at -0.003 g cm^-3^ (95%CI: -0.017 to 0.009), or -0.4% (-2.3 to 2.3, Table S2) of the predicted wood density. The mean uncertainty was 0.069 g cm^-3^ (or 12.4%, Fig. S2), and it varied substantially across space (95%CI: 0.049 to 0.087 g cm^-3^, or 8.0 to 16.5%).

To assess the importance of spatial autocorrelation, we tested how uncertainty depended on the distance of predictions from training samples (Table S3). Prediction errors were lowest directly adjacent to training data (0.043 g cm^-3^, i.e., 7.8% for a reference wood density of 0.55 g cm^-3^, Tables S4-5) but increased when training data were 100 km away (0.065 g cm^-3^, or 11.8%). Spatial autocorrelation reduced at larger distances, but uncertainty still increased up to 0.080 g cm^-3^ (14.5%) at 1,000 km. Random Forest models, our default approach, captured this spatial structure more effectively than linear regression models, but differences between approaches were small when modelled residuals were spatially interpolated (Fig. S3-4).

Uncertainties in community-weighted wood density were lowest in regions such as USA, which are densely sampled, with a mean uncertainty of 0.053 g cm^-3^ (10.4%, n_pixel_ = 7.0 million) and a 95% range from 0.048 to 0.067 g cm^-3^ (7.0 to 15.5%, locally down to 5%). By contrast, in South Africa, uncertainty reached 0.086 g cm^-3^ (12.8% of the mean, n_pixel_ = 0.8 million) and varied from 0.079 to 0.092 g cm^-3^ (11.2 to 13.7%). The highest relative uncertainties (up to 15-20%) were observed in the boreal zone where mean wood density was low and plot coverage was sparser than in the temperate zone (Table S2, Fig. 1b, Fig. S12). There was some evidence for regional prediction biases, such as underestimation at high and overestimation at low wood densities, but both were low, in the range of 0.01-0.02 g cm^-3^ (Fig. 1c and Fig. S5, Table S2).

### Environmental and biogeographic determinants of wood density

The most important predictor of community-weighted mean wood density variation was mean annual temperature (Fig. 2a, Fig. S6, Table 1). For an increase in temperature by one standard deviation (sd_MAT_ = 7.24°C), wood density increased by 0.075 g cm^-3^ (multiple regression model, 95% CI from 0.075 to 0.076), or roughly by 0.01 g cm^-3^ for a 1°C change. Site water balance was the next most important predictor (Fig. 2b, Table 1), with increasing water availability leading to a decrease in wood density (-0.012 g cm^-3^, 95% CI from -0.012 to -0.012, no change due to rounding, Fig. S6). These patterns were robust to substitution to other climatology layers (Table S1). Wood density further showed consistent, but moderate decreases of -0.017 g cm^-3^ with cyclone frequency (-0.017 to - 0.017) and gymnosperm proportion (-0.018 to -0.017), but their importance was low for wood density mapping (Table 1).

**Table 1.** Predictor importance and effect sizes for wood density mapping. Shown are 10 predictors used to create the global map of community-weighted mean wood density, their bivariate correlation with wood density (Pearson’s r), their effect size in a multiple regression model (g cm^-3^) and their importance in a random forest model (cf. methods). All predictors were standardized, so effect sizes correspond to shifts in wood density (in g cm^-3^) for a change of one standard deviation of the predictor variable. Predictors are sorted from most important to least important. This approach downweighs predictors with regional effects (e.g., cyclone frequency). Sources and definitions of the predictors can be found in Table S6. Correlations between predictors (Pearson’s r < 0.6 throughout) are shown in Figure S13.

| *Layer* | *Pearson's*<br>*r* | *Simple*<br>*regression*<br>*coefficient* | *Multiple*<br>*regression*<br>*coefficient* | *Importance*<br>*(spatial RF*<br>*validation)* |
| --- | --- | --- | --- | --- |
| Mean annual temperature ($^{\circ}\text{C}$) | 0.57 | 0.060 [0.060, 0.061] | 0.075 [0.075, 0.076] | 18 |
| Site water balance ($\text{kg m}^{-2} \text{ year}^{-1}$) | -0.22 | -0.024 [-0.024, -0.023] | -0.012 [-0.012, -0.012] | 5 |
| Soil sand content ($\text{g kg}^{-1}$) | -0.03 | -0.003 [-0.003, -0.003] | -0.003 [-0.003, -0.002] | 4 |
| Burned area (%) | 0.03 | 0.003 [0.003, 0.004] | -0.004 [-0.005, -0.004] | 3 |
| Deciduous angiosperms proportion (%) | -0.04 | -0.004 [-0.005, -0.004] | 0.015 [0.014, 0.015] | 3 |
| Elevation (m) | -0.17 | -0.019 [-0.019, -0.018] | 0.013 [0.013, 0.014] | 2 |
| Gymnosperm proportion (%) | -0.45 | -0.048 [-0.048, -0.047] | -0.017 [-0.018, -0.017] | 1 |
| Mean wind speed (m s<sup>-1</sup>) | -0.20 | -0.022 [-0.022, -0.021] | 0.011 [0.011, 0.012] | 1 |
| Cation exchange capacity (mmol(c) kg<sup>-1</sup>) | -0.33 | -0.035 [-0.035 -0.034] | 0.006 [0.006, 0.007] | 0 |
| Cyclone frequency (yr<sup>-1</sup>) | -0.04 | -0.005 [-0.005, -0.004] | -0.017 [-0.017, -0.017] | 0 |

**Fig. 2:**
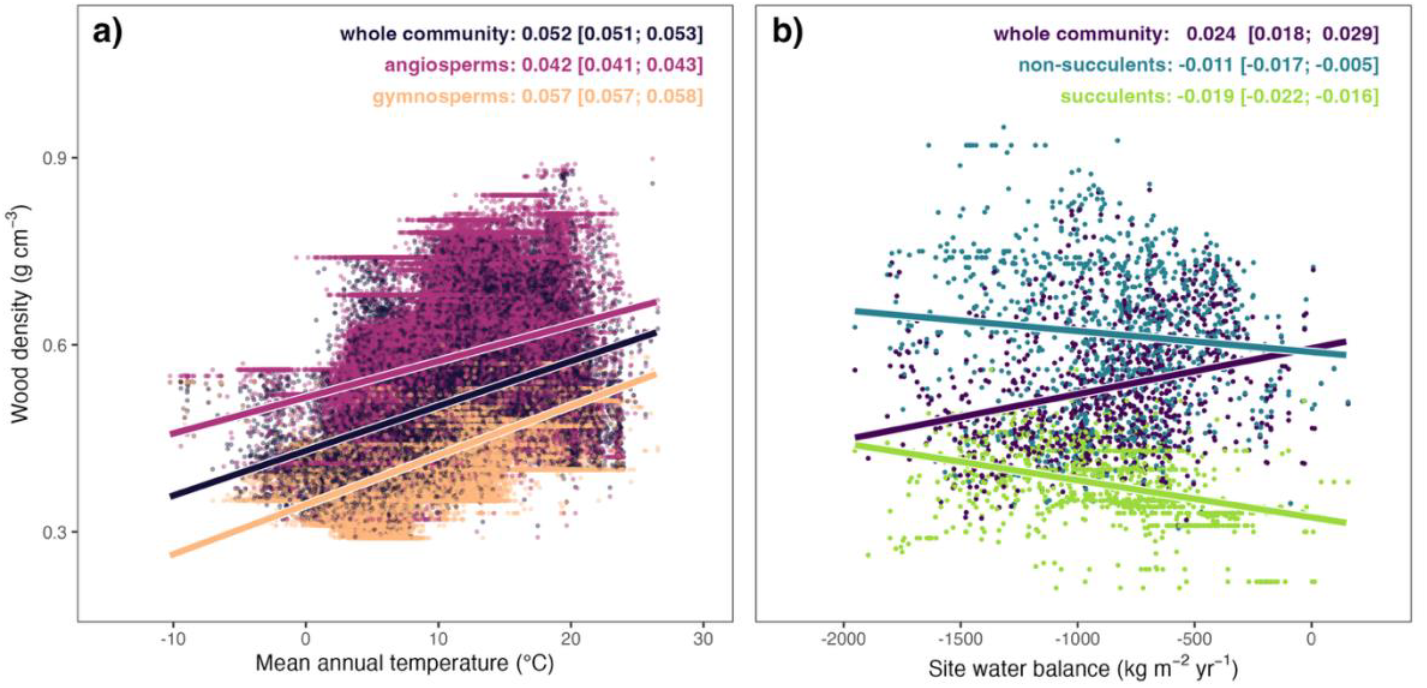
Examples of environmental wood density variation across phylogenetic groups and life forms. Panel a: relationship between mean annual temperature (°C) and community-weighted mean wood density (g cm^-3^) for plots where angiosperms and gymnosperms co-exist (n = 249,354). Wood density varies with temperature for angiosperms and gymnosperms separately as well as when both groups are combined (“whole community”). Lines are linear regression lines for each subset. The corresponding slope estimates and 95% CIs are provided in the top right corner. For comparability across predictors, slope estimates are reported with respect to standardized mean annual temperature, i.e., in units of g cm^-3^ for one standard deviation shift in temperature (sd_MAT_ = 7.24°C). Panel b: relationship between site water balance, an indicator of water availability for plant growth (sd_SWB_ = 320 kg m^-2^ yr^-1^, details in Table S6) and community-weighted mean wood density across communities with co-occurring succulents and non-succulents (n = 3,108). Succulents were defined broadly as trees or tree-like plants with any type of succulent habit (e.g., Baobab, Aloe tree, Saguaro). This includes stem-water storage and succulent roots or leaves. Note that wood density decreases with water availability in both subgroups in panel b, but increases when the subgroups are aggregated, which is due to decreasing succulent presence at high water availability (Simpson’s paradox). Such effects reduce predictive power when modelling community-weighted means (cf. Fig. S6b).

To assess the consistency and utility of temperature and aridity for wood density mapping, we computed temperature and aridity effects in two subgroups (Fig. 2), representing differences in xylem structure and leaf habit (angiosperms vs. gymnosperms) and drought resistance strategies (succulent vs. non-succulent woody plants). In mixed forests, wood density increased with temperature both at aggregate level (0.052 g cm^-3^, 95%CI: 0.051 to 0.053, simple linear regression) and separately in gymnosperms and angiosperms (0.057 g cm^-3^ and 0.042 g cm^-3^, with 95%CIs from 0.057 to 0.058, and from 0.041 to 0.043, respectively). Wood density also decreased with increasing water availability in co-occurring succulent (-0.019 g cm^-3^ for site water balance, -0.022 to -0.016) and non-succulent plants (-0.011 g cm^-3^, -0.017 to -0.005), but the declining presence of succulents in areas with higher water availability reversed the effect at aggregate levels (Fig. 2b).

### Comparison with alternative mapping approaches

We found that the present map was a significant improvement compared to previous mapping approaches. Uncertainty was reduced by 13% (ratio of variances) compared to a map produced with the previous GWDD (Fig. S7, uncertainty of 0.074 g cm^-3^, 95% interval from 0.49 to 0.99), and previous regional biases, such as an overestimation of wood densities across Australia (0.077 g cm^-3^ or 10.3% of mean wood density), were reduced to less than 2.5% (Fig. 1c). The present map also considerably improved over three published global wood density maps (Table S7), two of which calculated mean wood density directly from samples without weighting by basal area (“Boonman2020”^12^ and “Yang2024”^13^) and one that used the same definition of community-weighted means as our study (“Mo2024”^11^). There were similarities with the published maps in terms of global means (0.55 to 0.56 g cm^-3^) and qualitative trends (R^2^ of 0.60 to 0.83, Fig. S8-10, Table S7). However, our study revealed a 1.8-3.7 times larger spatial variation in wood density than published previously (Table S7), and we also improved mapping accuracy, reducing uncertainty by 51% compared to Boonman2020, 45% compared to Yang2024, and 32% compared to Mo2024.

Improvements were also noticeable regionally. Over Amazonian forests, we found a clear increase from 0.59 g cm^-3^ (95%CI: 0.55 to 0.63) in Southwest Amazon moist forests to 0.65 g cm^-3^ (95%CI: 0.61 to 0.69) in Guiana lowland moist forests (Fig 3a). This result replicated trends in the raw plot data (0.58 vs. 0.66 g cm^-3^) and a regional map (albeit weaker, Fig. S11). Overall, predictive power was low (R^2^ = 0.26), but comparable to the regional map (Fig. S11) and considerably higher than in previously published maps (R^2^ = 0.03-0.05), which predicted little to no variation across the Amazon (Fig. 3, b-d).

**Fig. 3:**
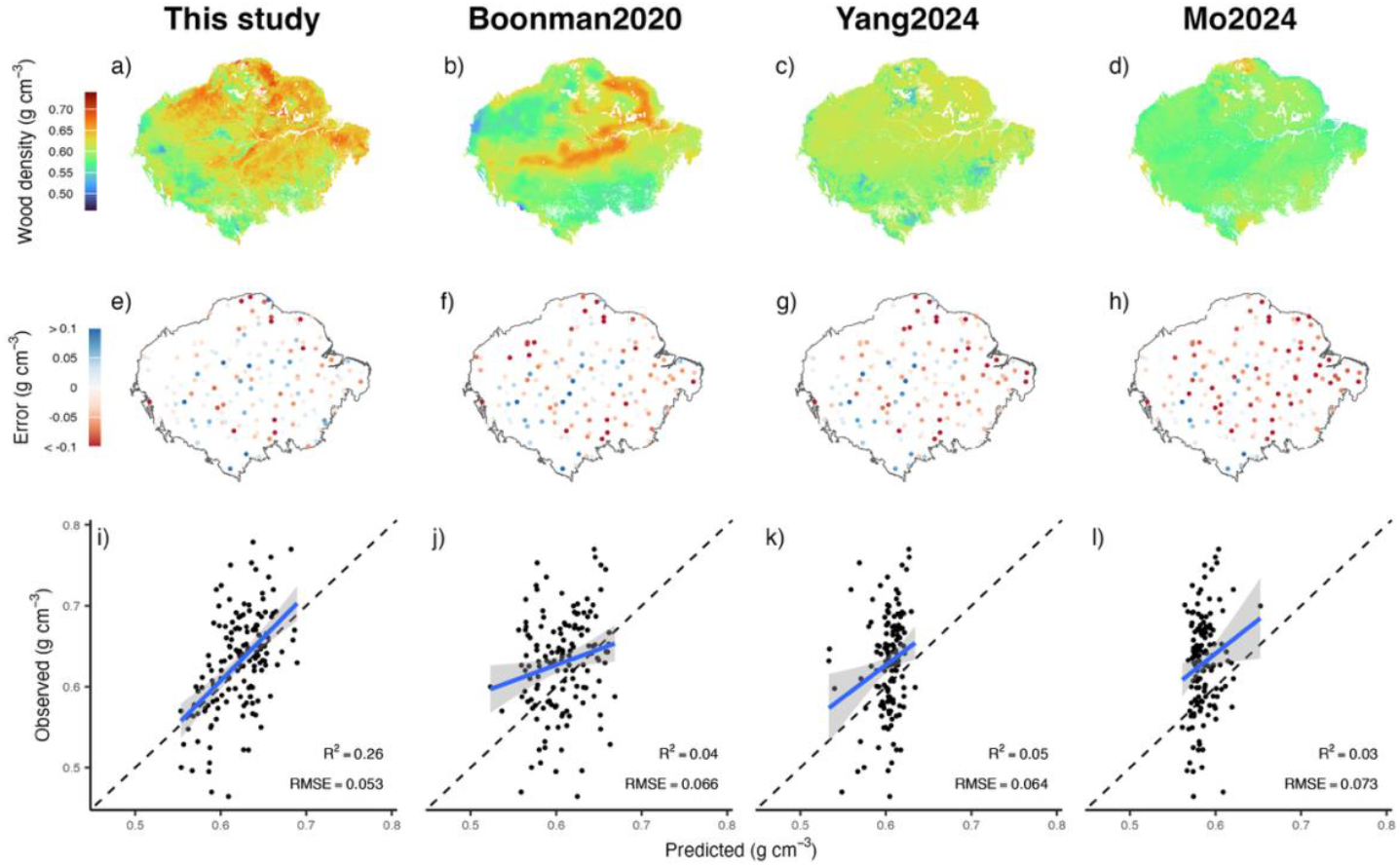
Wood density variation across Amazonian forests, as predicted from global wood density maps. Shown is the wood density gradient across Amazonia, as predicted by the present study and three published global maps (panels a-d), their errors across validation plots (e-h), and a plot of observed vs. predicted wood densities (i-l, n = 152, Fig. 1d). All depicted quantities are in g cm^-3^. Errors are computed via cross-validation (e and i) or direct map-plot comparisons (f-h and j-l). Blue lines in panels i-l are least squares regression lines with 95%CI envelopes. Note that variation in the validation data occurs at much smaller scales (∼1 ha plots) than the prediction scale (∼1 km^2^ pixels), which may explain why R^2^ values are low and observed wood density ranges larger in the validation data (approx. 0.45 to 0.8 g cm^-3^) than in any of the maps. For a comparison with a local wood density map, the same plot can be found in Fig. S11.

## Discussion

Here we mapped community-weighted mean wood density at global scale using a combination of wood density measurements and vegetation inventories, including nearly 17,000 species and more than 300,000 plot locations. The new wood density map achieved 32-51% higher accuracy than previous studies, revealed a 1.8-3.7 times larger variation and displayed clear environmental and biogeographic patterns. Our findings underscore the tight coupling between wood density and fundamental plant-ecological strategies^29^, which makes it a valuable predictive trait for modelling vegetation composition and dynamics across landscapes and through time^34^. We also carried out a detailed, spatial assessment of prediction errors, which revealed low biases (±2.5 % of the mean), and uncertainties ranging from 20% in the most poorly sampled regions to 5% in the best-sampled regions. Our work provides a critical improvement since wood density is a key component of carbon accounting at landscape^35,36^, regional^10^ and global scales^7,11^, requiring comprehensive, spatially explicit error budgets.

Our results differed from previous global studies that did not account for plant size when averaging at community-level^12,13^, but also outperformed an existing map of community-weighted mean wood density^11^ (Fig. 3, Figs. S8-10). Most of these discrepancies likely stemmed from the improved spatial coverage of plot collection in the present study and from the much larger and curated collection of wood density estimates of the GWDD v.2. By contrast, the inclusion of heavily human-modified forests in earlier training sets^11^ is unlikely to explain the narrower variance in those maps: strong anthropogenic disturbance would be expected to increase wood density variation among communities, not reduce it^37^. The performance and spatial patterns of the new map were more comparable to those seen in regional maps^15^, but there were still notable mismatches, for example in the strength and extent of gradients across Amazonia (Figure S11). To address these differences, future mapping efforts may benefit from considering additional soil and disturbance variables, which are expected to be the main drivers of wood density variation across Amazonia^14,15^.

The study at hand also showed that community-weighted mean wood density followed predictable patterns across large environmental gradients. Given balanced and high-quality inputs, most patterns could be predicted from a few biologically informative variables, potentially reducing the need for large predictor sets and complex model tuning^11,13,38^. For example, for an increase in temperature by one standard deviation, wood density was estimated to increase by 0.075 g cm^-3^, or by 0.01 g cm^-3^ for a 1°C change. Conversely, wood density decreased by -0.012 g cm^-3^ with a change in one standard deviation of site water balance, the second most important predictor. This confirmed the hypothesis that higher exposure to heat and drought stress favours slow-growing plants with dense wood^28^. There were regional deviations, e.g., high wood densities both in the driest (Sahel) and wettest parts (Congo Basin) of tropical Africa, but the strong and consistent relationship of community-level wood density with temperature was surprising (Fig. 2a). Temperature effects on wood density have rarely been explained physiologically^39^, and while they also exist at the species and intraspecific level, they usually remain less pronounced than the effects of water availability^33,38^. A possible explanation is that aridity tends to limit which strategies are viable in terms of building woody tissues, while temperature tends to determine which strategies are successful, e.g., at high aridity, plants with high wood density increasingly become viable, but, due to low growth rates, they do not dominate biomass accumulation until heat stress is also high. Global warming and increasing drought stress may thus lead to a reshaping of plant communities towards higher wood densities. To date, however, empirical studies do not support such a densification trend or even suggest the opposite pattern^40^.

More generally, the interpretation of environmental effects on wood density is complex. Climate variables such as temperature are linked to multiple stressors, and wood density is a compound trait that also summarizes several wood anatomical properties, including those of vessels, fibres and parenchyma cells^41^, and correlates with whole-plant architecture^42,43^. Links between wood density and the environment may thus arise indirectly^44^, or be mediated by tradeoffs^45^, trait-trait covariation^46^, and tree size^47^. Soluble organic compounds, so-called “extractives”, can also be a confounding factor, as they are usually not removed during wood density determination^48^. They have negligible impact on carbon estimates, as they account for a small share of stem biomass in most plants and possess carbon contents comparable to wood (40-50%)^49^. However, they may bias relationships between wood density and environmental factors and explain regional anomalies, such as the comparatively high wood densities in forests of extractive-rich larches^50^ across Siberia (Fig. 1a).

A key concern in mapping biological quantities across space is spatial autocorrelation, which comes about, for example, due to dispersal limitations of organisms, or unmodelled spatial variation in climatic and edaphic predictors, and which influences both model fitting^51^ and testing^31^. Here, prediction uncertainty in random forest models increased with distance from training data over the entire range that we tested, i.e., from 1 to 1,000 km, and most clearly (by 129%) up to 100 km. This finding reveals that a considerable portion of predictive power is due to spatial autocorrelation, far more than suggested by previous studies^11^. Confirmation of this result was provided by multiple regression models, which were generally inaccurate (R^2^ ∼ 0.3, Fig. S4), but improved substantially when spatially interpolated residuals were added to predictions (R^2^ up to 0.75). While spatial autocorrelation is often seen as a confounding factor in ecology, it also presents an opportunity to correct predictions through careful sampling protocols and spatial interpolation. For example, we estimated accuracy to be highest across systematically sampled regions such as USA (0.048 g cm^-3^, or down to 5%). Therefore, investment in gridded field data collections in poorly sampled regions (e.g., through national forest inventories and/or extension of plot networks) holds one of the keys to producing high-quality wood density and carbon maps^9^.

This study suggests two possible areas of improvement. Stressful environments spur a wide variety of strategies in plants to reduce mortality risks^52^. Wood density maps may considerably improve in accuracy if trade-offs between these strategies and wood density are better accounted for and more accurate predictor maps are created. In our study, succulents showed the same increase in wood density with aridity as co-occurring non-succulents (Fig. 2b). However, the generally light-wooded succulents increasingly dominated in dry areas – likely due to water storage and the often-co-occurring, transpiration-efficient CAM photosynthesis^53^ –, which reversed the relationship with aridity at aggregate levels (Simpson’s paradox, Fig. S6b). Similar patterns could emerge for other stress-resistance or stress-avoidance strategies (e.g., deciduousness), meaning reliable maps of stressors, life forms and growth habits are needed for wood density mapping.

Second, a large proportion of wood density variation occurs at scales smaller than 1 km^10^, reflecting the fine-scale mosaics of disturbance^37^ and species composition^19^. This scaling difference led to high uncertainties (low R^2^) when predicting wood density at plot-level from maps at km scales (Fig. 3). However, high-resolution forest age maps are now available^54,55^, and leveraging these maps together with high-resolution canopy heights maps^56^, would help map fine-scale wood density variation^36^. The present study will provide a robust foundation for future efforts and will help improve the accuracy of land carbon monitoring and vegetation modelling.

## Methods

### Wood density data assembly

We assembled an updated version of the Global Wood Density Database^32^, GWDD v.2^33^, which reports basic wood density, defined as oven-dried mass of a wood sample divided by its green volume^48^ in g cm^-3^. We employed the same wood density definition as in the original GWDD and, to maximize taxonomic coverage, included both measurements of individual plants and averages across several individuals. We used a broad definition of wood that includes tissues from tree-like plants that lack secondary growth, such as palms and succulents. We included direct measurements of basic wood density but also wood density values obtained at different moisture levels, which were transformed to basic wood density via physical conversion factors^18^. Taxonomic records were standardized via the *WorldFlora* package^57^ in R^58^, based on World Flora Online (www.worldfloraonline.org, June 2023 release), an international initiative that supersedes *The Plant List*^59^. Fuzzy matches of taxon names were manually checked and corrected where necessary. The final database contains 109,626 records from 16,829 species, with coverage across biomes and biogeographic realms^33^.

### Plot data assembly

To weight wood density by relative species dominance, we assembled a global collection of 300,949 vegetation inventories – plots with taxonomic identification and measurements of stem diameter for all sampled plants – from 300,702 unique 1 km x 1 km cells and 281 sources (Table S8, Fig. S12). We privileged old-growth and intact forests and, where identifiable, excluded data from woody ecosystems with strong anthropogenic impacts. We selected primarily ecological studies from protected areas, excluded heavly managed stands from national forest inventories and plantations of non-native species, such as eucalypts planted outside of Australia. Woody ecosystems such as xeric environments with succulents or shrubs were included, with no minimum threshold in woody cover, to represent the full range of wood density variation.

For each plot, we defined the community-weighted wood density CWWD by averaging across individuals i as follows:

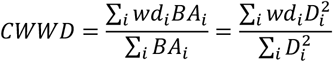

Here *wd_i_* is the wood density of individual i, 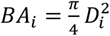 its cross-sectional (or basal) area and *D_i_* its trunk diameter. This weighting scheme gives considerably more weight to species with large canopy trees than a simple aggregation across all individuals (i.e., weighting by species abundance), but less than weighting by tree volume, since the latter depends not only on basal area, but also tree height, which also increases with trunk diameter^60^. While weighting schemes subtly shift the interpretation of the typical traits of communities, in practice, community-weighted means of wood density have proven robust to how different size classes are weighted^15,31^, and basal-area weighting remains a widely used and practical approach^19^.

The majority of the CWWD values were computed from open access raw data on tree- or species level basal area (n = 243,776). We also identified geographical data gaps– primarily dry ecosystems in South America, South and East Africa, India –, and supplemented the collection with published surveys from these areas. Additional geographic gaps were filled by direct CWWD estimates from the literature, especially in tropical forests^10,31,61^ and in Russian forests^62^. Some of these published CWWD values have been computed from the original GWDD, so they may carry over some biases due to conversion factors^18^, but they fill an important knowledge gap and were therefore retained. Most of the surveys included trees measured at 10 cm in trunk diameter and above, but this threshold varied from 1 up to 30 cm in some locations. This should not systematically bias wood density estimates, since community-weighted means tend to be robust to the choice of weighting factors^15^ and since large trees disproportionally contribute to basal area and woody volume^31^.

Inventory data were matched to wood density values from the GWDD v.2 at the species and genus level following standard procedures^63^. Only plots with wood density values for >80% of the total basal area were retained. If two or more plots from the same source fell into the same 1 km x 1 km grid cell (Mollweide equal-area projection, ESRI:54009), their CWWD values were averaged to produce a cell-wide CWWD estimate using the *terra* package^64^. To do this, plots were averaged by weighting by their total basal area.

### Wood density mapping

To map wood density, we assembled 10 climatic, disturbance and auxiliary layers (full definitions in Table S6), and resampled them to the 1 km x 1 km Mollweide grid also used for aggregating wood density estimates. We chose predictors in accordance with biological hypotheses about the determinants of wood density variation (stress, disturbance, soil fertility), including climatic variables for the period 1981-2010 from CHELSA/BIOCLIM+^65,66^, edaphic factors^67^, disturbance agents such as fire^68^ and broad-scale community composition^69^. We also derived cyclone frequency from the best track archive^70^ by counting how often each 1 km x 1 km grid cell was within a 150 km radius of a cyclone track and normalizing this number by the number of years. Pairwise correlations between predictors never reached r = 0.6, which minimizes collinearity issues with attributing effects when modelling and mapping wood density^71^.

Based on the set of 10 predictors, we applied random forest models to predict wood density at 1 km x 1 km resolution. We used the *ranger* package^72^. To account for spatial autocorrelation in wood density variation and improve predictions, we added spatially interpolated residuals to the final map, relying on the 25 nearest plots for each grid cell and using inverse distance weighting (1/r, where r is a plot’s distance from the target grid cell) via the R packages *terra* and *gstat^73^*. For comparison, we also retained the map without spatially interpolated residuals. We repeated the procedure with a multiple regression model with the same 10 predictors, again creating maps both with and without spatially interpolated residuals. We mapped wood density across all land pixels globe but then masked pixels without any woody plants (0% tree or shrub cover), using a mask from the Copernicus Global Land Cover product^74^.

### Map validation

We validated the map using a subset of 1,000 plots, selected as follows. We chose a minimum sampling area of 0.25 ha to reduce noise, then selected plots via geographic stratification to avoid biasing validation towards regions of denser sampling. To this end, we subdivided the globe into 200 x 200 km grid cells, and randomly sampled one plot from each grid cell. After having drawn once from each grid cell, continued drawing from random grid cells until reaching 1,000 samples (Fig. S1). We selected 1,000 plots, as this reflects well the distance between plots in sparsely sampled areas (∼100-200 km) and provides a large enough sample size to reduce randomness in validation.

Once the validation plots were selected, we carried out a spatial leave-one-out (LOO) cross validation^31^. First, we carried out a standard LOO validation: for each of the 1,000 plots, we replicated the wood density map, including spatial interpolation of residuals, based on all data except the validation plot itself and compared predicted against observed wood density values. We then repeated this process three times, but each time removed not only the validation plot from the training data, but also all data points within an exclusion radius ranging between 1 and 1,000 km from the validation plot.^30^ We did not choose the exclusion radii as fixed thresholds (as in ref. ^31^), but drew them at random from a uniform distribution. This choice allowed us to assess spatial autocorrelation continuously up to 1,000 km. A few duplicates occurred when randomly drawn exclusion radii did not exclude any additional plots. We removed these predictions, resulting in a total of 3,841 data points for validation.

To generate error estimates for each grid cell in the wood density map, we modelled the error distribution depending on the predicted wood density, the distance from the nearest five plots and the interaction of both (Table S3). The best fit to the data was obtained by modelling uncertainty with a *t* location-scale or “student” distribution, which is useful for modeling data distributions with heavier tails than the normal distribution:

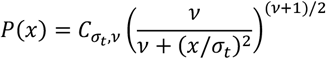

This distribution approaches the normal distribution as *v* approaches infinity, while smaller values of *v* yield heavier tails. With this distribution, the variance is given by *var* = σ × *v*/(*v* − 2). The fitting was performed with the package *brms^75^* and suggested a shape parameter of either *v* = 5 or *v* = 6. To ensure comparable error structure across models, we fixed *v* = 5 and then mapped both the mean error (bias) and uncertainty in wood density. Uncertainty can be directly expressed via the σ_t_ parameter, but here we report 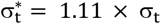, which is comparable to the standard deviation of a normal distribution, since 68.26% of data points lie within 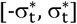.

### Assessment of environmental determinants

To evaluate the relative importance of environmental variables for wood density mapping we re-created the global wood density map 10 times, each time dropping one of the predictors, carrying out a spatial cross-validation and mapping uncertainty across woody ecosystems. Predictors that, upon dropping, led to the largest loss of accuracy globally, were considered the most important for the mapping. To reduce the computational burden in spatial cross-validation, we created a reduced data set of n = 74,725 plots by applying a 10 km x 10 km filter and selecting a single plot per grid cell for model training. We verified that this choice had negligible impacts on model accuracy. For an assessment of the direction of effects, we also computed simple linear regression models, with community-weighted mean wood density regressed against each predictor separately, and fitted a multiple linear regression model with the same predictors. More precisely, we used the model 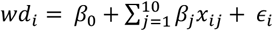, where *wd_i_* is the community weighted wood density of plot *i, x_ij_* is predictor variable *j* for observation *i*, and ϵ*_i_* is the residual for observation *i*, distributed as ϵ*_i_*∼ *N*(0, σ^2)^. Predictors were scaled to one standard deviation to ensure comparability of effect sizes.

Climatic predictors were found to be the most important ones in predicting spatial variation in wood density, so we decided to carry out two additional robustness tests. First, we used the TerraClimate^76^ and ERA5-Land^77^ datasets in place of the CHELSA data and evaluated changes in the global wood density map. In the case of mean annual temperature and mean wind speed, we used like-for-like substitutions. In the case of CHELSA’s site water balance variable, there was no direct equivalent, so we chose other aridity-related predictors, such as climatic water deficit, the difference between potential and actual evapotranspiration (TerraClimate), and potential evapotranspiration (ERA5-Land).

Variation in wood density was also found to depend on wood anatomy and plant life forms. If either of those factors changed the effect of environmental predictors, not or insufficiently accounting for them could induce bias and increase uncertainty in wood density maps. We tested the consistency of predictions across gymnosperms and angiosperms, as these are species-rich and globally dominant plant groups that differ in wood anatomy and leaf habit, and across succulents and non-succulents, as water storage in plant organs is a key drought-resistance strategy, which has often co-evolved with transpiration-reducing CAM photosynthesis^53^ and should lead to strong trade-offs with wood density in dry regions^52^. For the former, we selected all plots with both angiosperms and gymnosperms (n = 249,354, “mixed forests”) and compared simple linear regression estimates of community-weighted wood density against temperature within each plant group and across groups. For the latter, we matched diameter measurements to a list of succulents compiled from the literature^78–81^ and compared regression estimates of community-weighted wood density against site water balance both within each life form and across life forms. The full list of succulent species with wood density records in the GWDD v.2 can be found in Table S8. We did not distinguish between different types of succulence, and defined succulents broadly as woody or wood-like plants with any type of succulent habit.

### Comparison with wood density maps

Finally, we compared the global wood density map to other products. First, we compared it to a map created only from wood density records in the previous version of the GWDD^32^. We compared both maps in terms of overall uncertainty and assessed regional biases due to lower species coverage and outdated conversion factors^18^. Second, we compared it to three published maps^11–13^. Although two of these maps did not weight wood density by tree size (basal area or volume), their estimates were explicitly aimed at representing communities in terms of species dominance and carbon stocks^12,13^, and should thus be comparable with the community-weighted means used in this study. We resampled all three maps to the same 1 km x 1 km Mollweide grid, then evaluated them in terms of overall wood density distribution and against the same 1,000 spatially stratified validation plots. We assessed correlations with the reference map via R^2^, and predictive performance on validation plots via root mean square errors (RMSE) and R^2^. The latter were compared to the RMSE and R^2^ of the reference map under basic leave-one-out cross validation (ratio of variances). Third, we assessed the ability of our reference map and the three published maps to predict a known wood density gradient across Amazonian rainforests^10^. We compared RMSEs and R^2^ across plots from the global validation dataset (Fig. 1d), but located within Amazonia (n = 152), and also compared predictions from this study against a recently produced regional map ^15^ (Fig. S11).

## Acknowledgments

The production of global-scale trait and carbon maps would not be possible without the diligent work of a large scientific community and the many openly available datasets and tools that we relied on here. We acknowledge the work of thousands of trait collectors and wood technicians; taxonomists who maintain and harmonize species records; foresters, ecologists and field assistants who measure trees on the ground; meteorologists who operate weather stations and collect climate data; satellite engineers, climate scientists and IT developers who develop models and maps as well as the software to analyze them; as well as individual citizens, local communities, and public and private bodies that support this work.

F. J. Fischer acknowledges a European Research Council grant to Rupert Seidl under the European Union’s Horizon 2020 research and innovation program (Grant Agreement 101001905, FORWARD). J. Chave acknowledges an ‘Investissement d’Avenir’ grant managed by the Agence Nationale de la Recherche (CEBA grant: ANR-10-LABX-25-01 and TULIP grant: ANR-10-LABX-0041). T. Jucker acknowledges a UK NERC Independent Research Fellowship (grant: NE/S01537X/1), UKRI Frontier Research grant (grant: EP/Y003810/1) and a Research Project Grant from the Leverhulme Trust (grant: RPG-2020-341). A. Fajardo acknowledges Anid-Fondecyt 1231025. R.A.F. de Lima acknowledges grant 13/08722-5, São Paulo Research Foundation (FAPESP). G. Vieilledent acknowledges the French Foundation for Research on Biodiversity (BIOSCENEMADA project). L. F. Alves acknowledges a Smithsonian Tropical Research Institution, Center for Tropical Forest Science (CTFS) Research Grant (2009). J. Borah acknowledges the village councils of Kiphire (Fakim, Thanamir, and Tsundang village), Phek (Zhipu, Wazeho, and Washelo village), and Kohima (Dzuleke village) district, Nagaland, India. J. Camarero acknowledges the Spanish Ministry of Science and Innovation (projects PID2021-123675OB-C43 and TED2021-129770B-C21). E. Cifuentes acknowledges Convocatoria 860 MinCiencias, Colombia. J.A. Dar acknowledges the Anusandhan National Research Foundation (ANRF), Government of India, for funding (Ref. No.: SRG/2022/002286) and SRM University-AP for the Seed Grant (SRMAP/URG/E&PP/2022–23/012). A. K. Das acknowledges a research project grant from the Council of Scientific and Industrial Research, New Delhi (Government of India, project no.38(1349)/13/EMR-II dt.14.2.2013). P. Fearnside acknowledges the National Council for Scientific and Technological Development (CNPq 12450/2021-4, 406941/2022-0), the Foundation for the Support of Research in the State of Amazonas (FAPEAM: 01.02.016301.02529/2024-87) and the Brazilian Research Network on Climate Change (RedeClima) (FINEP/Rede Clima 01.13.0353-00). S. Kothandaraman acknowledges the Science and Engineering Research Board, Department of Science and Technology (DST-SERB) (Ref. No.: PDF/2021/003742/LS). A. Linstädter, M. Dobler and L. Kindermann acknowledge funding by German Research Foundation (DFG) through funding codes CRC TRR-228/1 and TRR-228/2, and thank the University of Namibia (UNAM) as well as National Botanical Research Institute Windhoek for their support. A.L.A Lima acknowledges the National Council for Scientific and Technological Development (CNPq). Pernambuco Science and Technology Support Foundation (FACEPE). Cate Macinnis-Ng acknowledges a Rutherford Discovery Fellowship from the Royal Society Te Apārangi RDF-UOA1504; Marsden Fund Award from the Royal Society Te Apārangi UOA1207. L.F.S. Magnago acknowledges the Conselho Nacional de Desenvolvimento Científico e Tecnológico (CNPq), grant 307984/2022-2. A. Martin acknowledges the Natural Sciences and Engineering Research Council of Canada. A. Matheny acknowledges an NSF EAR Hydrological Sciences CAREER Award 2046768. E. Nogueira acknowledges the National Institute for Research in Amazonia (INPA). O. Razafindratsima acknowledges the Madagascar National Parks and the Malagasy Ministry of Environment for research permission, and numerous local field workers. S. Richardson acknowledges the Strategic Science Investment Fund administed by the NZ Ministry for Business, Innovation and Employment. Oris Rodriguez-Reyes acknowledges the Instituto de Ciencias Ambientales y Biodiversidad, Universidad de Panama; Sistema Nacional de Investigación Panama; STRI Panama. R. Salguero-Gomez acknowledges a NERC Independent Research Fellowship (NE/M018458/1) and a NERC Pushing the Frontiers grant (NE/X013766/1). L. F. Henao-Diaz and P. Stevenson acknowledge Ecopetrol No. 135-2009. M. van der Sanda acknowledges a Veni grant from the Dutch Research Council (NWO), NWO-VI.Veni.192.027. J.M.D. Torezan acknowledges a grant from the Brazilian science council (CNPq PELD 441510/2020-5). In addition, we thank Jordi Martínez-Vilalta and Teresa Rosas for their data contributions, supported by CGL2013-46808-R grant funded by MINECO, and Dmitry Schepaschenko for his contributions to the data assembly and facilitating the coordination among contributors.

## Author contributions

F.J. Fischer, J. Chave and A. Zanne conceived of the project. F.J. Fischer led the assembly and processing of the data, the analysis for the manuscript, and the writing of a first draft of the manuscript, with assistance from J. Chave, A. Zanne, T. Jucker, A. Fajardo, A. Fayolle, R.A.F. de Lima and G. Vieilledent. All co-authors contributed substantially through data collection, data assembly and revisions of the manuscript.

## Data availability

All data sources for this analysis are openly available. The GWDD v.2 will be made publicly available on Zenodo (doi: 10.5281/zenodo.16919510) as part of a companion manuscript. All code to reproduce the analyses and maps in this manuscript will be made publicly available on Zenodo upon publication of this manuscript (doi: 10.5281/zenodo.16932871). The references for all data sources and the analysis scripts are available in the Supplementary Information.

