## Supplementary Materials for "A global map of wood density"

### A. Supplementary Tables

| *Biome* | *CHELSA /BIOCLIM+* | *TerraClimate* | *ERA5-Land* |
|---|---|---|---|
| Boreal Forests/Taiga | 0.44 | 0.44 | 0.44 |
| Deserts & Xeric Shrublands | 0.66 | 0.66 | 0.65 |
| Flooded Grasslands & Savannas | 0.62 | 0.62 | 0.61 |
| Mangroves | 0.62 | 0.61 | 0.62 |
| Mediterranean Forests, Woodlands & Scrub | 0.65 | 0.65 | 0.65 |
| Montane Grasslands & Shrublands | 0.52 | 0.52 | 0.52 |
| Temperate Broadleaf & Mixed Forests | 0.51 | 0.51 | 0.51 |
| Temperate Conifer Forests | 0.45 | 0.45 | 0.45 |
| Temperate Grasslands, Savannas & Shrublands | 0.57 | 0.57 | 0.56 |
| Tropical & Subtropical Coniferous Forests | 0.61 | 0.60 | 0.61 |
| Tropical & Subtropical Dry Broadleaf Forests | 0.64 | 0.62 | 0.63 |
| Tropical & Subtropical Grasslands, Savannas & Shrublands | 0.66 | 0.66 | 0.66 |
| Tropical & Subtropical Moist Broadleaf Forests | 0.60 | 0.60 | 0.61 |
| Tundra | 0.46 | 0.45 | 0.45 |

**Table S1. Wood density by biome, for three different sources of climatic predictors.** Shown are the average community-weighted wood densities by biome (with boundaries according to Dinerstein et al., 2017) for the default map where the climatic predictors are derived from CHELSA/BIOCLIM+ (Brun et al., 2022; Karger et al., 2017), as well as alternative maps that use comparable predictors from two different climate products, TerraClimate (Abatzoglou et al., 2018) and ERA5-Land (Muñoz-Sabater et al., 2021). Note that there are 10 predictors in total, and climatic predictors constitute only a subset of 3 predictors, but two of them (temperature and aridity) are the most important ones by far.

| *Biome* | *Uncertainty (g cm$^{-3}$)* | *Relative. Uncertainty (%)* | *Bias (g cm$^{-3}$)* | *Relative bias (%)* |
|---|---|---|---|---|
| Boreal Forests/Taiga | 0.066 | 15 | 0.006 | 2 |
| Deserts & Xeric Shrublands | 0.073 | 11 | -0.010 | -1 |
| Flooded Grasslands & Savannas | 0.074 | 12 | -0.008 | -1 |
| Mangroves | 0.072 | 12 | -0.007 | -1 |
| Mediterranean Forests, Woodlands & Scrub | 0.069 | 11 | -0.009 | -1 |
| Montane Grasslands & Shrublands | 0.070 | 14 | 0.000 | 0 |
| Temperate Broadleaf & Mixed Forests | 0.064 | 13 | 0.001 | 0 |
| Temperate Conifer Forests | 0.059 | 13 | 0.004 | 1 |
| Temperate Grasslands, Savannas & Shrublands | 0.064 | 12 | -0.003 | 0 |
| Tropical & Subtropical Coniferous Forests | 0.057 | 9 | -0.005 | -1 |
| Tropical & Subtropical Dry Broadleaf Forests | 0.072 | 11 | -0.008 | -1 |
| Tropical & Subtropical Grasslands, Savannas & Shrublands | 0.077 | 12 | -0.010 | -2 |
| Tropical & Subtropical Moist Broadleaf Forests | 0.068 | 11 | -0.006 | -1 |
| Tundra | 0.068 | 15 | 0.005 | 1 |

**Table S2. Wood errors by biome.** Shown are both uncertainty and bias for average community-weighted wood densities by biome (with boundaries according to Dinerstein et al., 2017) for the default map where the climatic predictors are derived from CHELSA/BIOCLIM+ (Brun et al., 2022; Karger et al., 2017). Relative values are calculated as percentages of the mean wood density.

| | Wood density error (in g cm$^{-3}$) | |
|---|---|---|
| *Predictors* | *Estimates* | *CI (95%)* |
| intercept | 0.02 | -0.02 – 0.06 |
| log-distance | 0.00 | -0.00 – 0.01 |
| wd | -0.03 | -0.10 – 0.03 |
| log-distance:wd | -0.01 | -0.02 – 0.00 |
| intercept$_{\sigma}$ | -2.91 | -3.65 – -2.17 |
| log-distance$_{\sigma}$ | -0.03 | -0.16 – 0.11 |
| wd$_{\sigma}$ | -0.63 | -1.85 – 0.63 |
| log-distance$_{\sigma}$:wd$_{\sigma}$ | 0.21 | -0.01 – 0.43 |
| Observations | 3841 | |

**Table S3. Model to predict errors in wood density map.** Shown are results from a model that predicts both the mean error (bias) and the spread of errors around the predicted mean (uncertainty, $\sigma_t$) as a function of the log-transformed distance to the five nearest training samples (log-distance), their community-weighted wood density (wd), and the interaction term between the two. The model was fit with the brms package (Bürkner, 2018), using standard formulae (error ~ log-distance + wd + log-distance * wd; sigmaerror ~ log-distance + wd + log-distance * wd; where sigmaerror is modelled on exponential scales), a student response distribution with 5 degrees of freedom and 1,000 validation plots. Each plot is included four times, once through a simple leave-one-out cross validation, and an additional three times with a random exclusion radius (up to 1,000 km) from within which training data were removed in a spatial cross validation approach (Ploton et al., 2020). For the final model, we included only validation plots which had their five nearest training samples within a 1000 km radius to avoid excessive influence of a few outlier points, and we removed any duplicates that occurred when the exclusion radius was smaller than the original data points distance to the nearest training data. Hence, we only used 3,841 instead of a potential 4,000 validation data points overall. For more information on "distributional models" cf. https://cran.r-project.org/web/packages/brms/vignettes/brms_distreg.html.

| *Distance to five nearest neighbours* | Prediction uncertainty ($\sigma_t$, in g cm$^{-3}$) at three wood density levels | | |
|---|---|---|---|
| | ***0.45 g cm$^{-3}$*** | ***0.55 g cm$^{-3}$*** | ***0.65 g cm$^{-3}$*** |
| 1 | 0.041 | 0.039 | 0.036 |
| 10 | 0.048 | 0.048 | 0.047 |
| 20 | 0.051 | 0.051 | 0.051 |
| 50 | 0.054 | 0.055 | 0.056 |
| 100 | 0.057 | 0.059 | 0.061 |
| 200 | 0.059 | 0.062 | 0.066 |
| 500 | 0.063 | 0.068 | 0.073 |
| 1000 | 0.066 | 0.072 | 0.079 |

**Table S4. Predicted errors in wood density estimates, original variance parameter.** Shown is the predicted uncertainty in wood density estimates (model in Table S1) at various distances from the nearest five training data points and for various levels of wood density. Uncertainty is described by the $\sigma_t$ parameter of a student distribution with 5 degrees of freedom. Cf. Table S3 for a transformation of $\sigma_t$ into $\sigma_t$*, which yields values comparable to the standard deviation of a Gaussian distribution.

| *Distance to five nearest neighbours* | **Prediction uncertainty ($\sigma_t$*, in g cm$^{-3}$) at three wood density levels** | | |
|---|---|---|---|
| | ***0.45 g cm$^{-3}$*** | ***0.55 g cm$^{-3}$*** | ***0.65 g cm$^{-3}$*** |
| 1 | 0.046 (10%) | 0.043 (8%) | 0.040 (6%) |
| 10 | 0.053 (12%) | 0.053 (10%) | 0.052 (8%) |
| 20 | 0.057 (13%) | 0.057 (10%) | 0.057 (9%) |
| 50 | 0.060 (13%) | 0.061 (11%) | 0.062 (10%) |
| 100 | 0.063 (14%) | 0.065 (12%) | 0.068 (10%) |
| 200 | 0.065 (15%) | 0.069 (13%) | 0.073 (11%) |
| 500 | 0.070 (16%) | 0.075 (14%) | 0.081 (12%) |
| 1000 | 0.073 (16%) | 0.080 (15%) | 0.088 (13%) |

**Table S5. Predicted errors in wood density estimates, transformed to Gaussian-like errors.** Shown is the predicted uncertainty in wood density estimates (model in Table S1) at various distances from the nearest five training data points and for various levels of wood density. This is the same table as Table S2, but uncertainty is here described by $\sigma_t$*, which is defined as $\sigma_t^* = 1.11 \times \sigma_t$, with $\sigma_t$ being the variance parameter of a student distribution with 5 degrees of freedom. Approximately 68.3% of values in the student distribution will lie within [-$\sigma_t$*,$\sigma_t$*], so $\sigma_t$* can be thought of as an equivalent of the standard deviation in Gaussian error models (or related error quantities such as RMSE). In addition, percentage errors are provided in brackets.

| *Layer* | *Definition* | *Source* |
|---|---|---|
| Mean annual temperature (°C) | Mean annual daily mean air temperatures averaged over one year (also “bio1”). | (Karger et al., 2017; Brun et al., 2022) |
| Site water balance (kg $m^{-2}$ $year^{-1}$) | Estimate of the water available to plants during a year that considers soil parameters in addition to climate variables. Its maximum is given by available water holding capacity of the soil. Minimum values indicate that evapotranspiration has exceeded precipitation minus runoff (also “swb”, cf. documentation at https://chelsa-climate.org/bioclim/) | (Karger et al., 2017; Brun et al., 2022) |
| Soil sand content (g $kg^{-1}$) | Gravimetric content of sand in the fine earth fraction of the soil in the standard depth interval 0-5 cm. | (Hengl et al., 2017; Poggio et al., 2021) |
| Burned area (%) | Percentage of area that is burned annually. | (Giglio et al., 2018) |
| Deciduous angiosperm proportion (%) | Proportion of land covered by deciduous angiosperms. | (Tuanmu & Jetz, 2014) |
| Elevation (m) | Elevation above mean sea level in meters relative to the Earth Gravitational Model of 2008 (EGM2008) geoid surface (EPSG:3855). | (NOAA National Centers for Environmental Information, 2022) |
| Gymnosperm proportion (%) | Proportion of land covered by gymnosperms. | (Tuanmu & Jetz, 2014) |
| Mean wind speed (m $s^{-1}$) | Average monthly near-surface wind speed over one year; near surface represents 10 m above ground (also “sfcWind_mean”). | (Karger et al., 2017; Brun et al., 2022) |
| Cation exchange capacity (mmol(c) $kg^{-1}$) | Cation Exchange Capacity of the soil, estimate of cation retention in soil. | (Hengl et al., 2017) |
| Cyclone frequency ($yr^{-1}$) | Average yearly number of passages of cyclones within 150 km. | (Knapp et al., 2010) |

**Table S6. Predictors and their sources.** Shown are the 10 predictors used for wood density mapping, their definitions as well as the references from which they were derived. In most cases, definitions are taken verbatim or in slightly altered form from the references. Note that Tuanmu & Jetz, 2014 provide a wide range of land classes, including both evergreen angiosperm proportion and several non-forest land classes, so the two predictor layers “deciduous angiosperm proportion” and “gymnosperm proportion” do not sum to 1.

| | Reference map | Boonman2020 | Yang2024 | Mo2024 |
|---|---|---|---|---|
| Mean wood density (g $cm^{-3}$) | 0.57 | 0.56 | 0.56 | 0.55 |
| Standard deviation (g $cm^{-3}$) | 0.104 | 0.054 | 0.059 | 0.074 |
| Ratio of variances (reference / published map) | (1.0) | 3.7 | 2.9 | 1.8 |
| $R^2$ (reference vs. published map) | (1.0) | 0.60 | 0.61 | 0.83 |
| RMSE (g $cm^{-3}$, validation plots vs. map) | 0.074 | 0.106 | 0.100 | 0.090 |
| $R^2$ (validation plots vs. map) | 0.69 | 0.38 | 0.42 | 0.50 |
| $n_{validation}$ | 998 | 934 | 776 | 828 |

**Table S7. Comparison of global wood density maps.** Shown is a comparison of the global wood density map created in this study ("reference map") with three published maps (cf. description in main text). All maps use different definitions of woody vegetation extent, so comparisons rely on pairwise complete observations, i.e., for each summary statistic (mean, standard deviation, etc.), we used only pixels with valid predictions both in the published and the reference map. The ratio of variances refers to the ratio of wood density variance in the reference map, divided by the variance in the published map. The last three rows represent the results from a validation of all four maps against a set of 1,000 geographically balanced validation plots. Again, estimates of woody cover differ between maps, so some validation plots do not have valid predictions, and $n_{validation}$ provides the actual number of validation plots for each map. For the map in this study, we estimated RMSE and $R^2$ from basic cross-validation, for the other maps we directly compared observed values with map predictions. Note that this may overestimate predictive power of the published maps, since some of the validation plots may have been used in their creation.

| ***plot type*** | ***source*** | ***median area (ha)*** | ***#plots*** |
|---|---|---|---|
| NFI | FIA. https://www_fia.fs.fed.us/ | 0.07 | 153498 |
| NFI | NFI Mexico. Second cycle. https://datos.gob.mx | 0.64 | 20408 |
| NFI | NFI Spain. Third cycle. https://www.miteco.gob.es | 0.20 | 58175 |
| NFI | NFI/CMI British Columbia, https://www2.gov.bc.ca/gov/content/industry/forestry/managing-our-forest-resources/forest-inventory/national-forest-inventory-program | 0.04 | 227 |
| NFI | NFI/PEP Quebec, https://www.donneesquebec.ca/recherche/fr/dataset/placettes-echantillons-permanentes-1970-a-aujourd-hui | 0.06 | 4251 |
| NFI | RADAMBRASIL | 1.00 | 2945 |
| phytosoc. study | Kartawinata et al. 2004. A tree species inventory in a one-hectare plot at the Batang Gadis National Park, North Sumatra, Indonesia. Reinwardtia 12, 145-157. | 1.00 | 1 |
| phytosoc. study | TERN (2022) AusPlots ecosystem surveillance monitoring dataset (URL: http://aekos.org.au/). Obtained via the ausplotsR R package (URL: https://github.com/ternaustralia/ausplotsR), accessed 29 June 2022. | 1.00 | 382 |
| phytosoc. study | Phumphuang et al. 2024: Environmental factors differentially influence species distributions across tree size classes in a dry evergreen forest in Sakaerat Biosphere Reserve, northeastern Thailand, Journal of Forest Research, DOI: 10.1080/13416979.2024.2314834 | 16.00 | 1 |
| phytosoc. study | Moreira et al. 2021. Species composition and diversity of woody communities along an elevational gradient in tropical Dwarf Cloud Forest. Journal of Mountain Science 18(6). https://doi.org/10.1007/s11629-020-6055-x | 1.00 | 1 |
| phytosoc. study | França & Stehmann 2004. Composição florística e estrutura do componente arbóreo de uma floresta altimontana no município de Camanducaia, Minas Gerais, Brasil. Brazilian Journal of Botany, 19-30. | 0.75 | 1 |
| phytosoc. study | Culmsee et al. 2011. Tree diversity and phytogeographical patterns of tropical high mountain rain forests in Central Sulawesi, Indonesia. Biodivers Conserv 20, 1103-1123. | 1.92 | 2 |
| phytosoc. study | Diogo et al. 2021. Effects of topography and climate on Neotropical mountain forests structure in the semiarid region. Appl Veg Sci, 24:e12527. https://doi.org/10.1111/avsc.12527 | 3.00 | 1 |
| phytosoc. study | Dohn et al. 2017. Spatial vegetation patterns and neighborhood competition among woody plants in an East African savanna. Ecology 98, p. 478-488. | 0.75 | 1 |
| phytosoc. study | Paul Musili Mutuku & David Kenfack 2019. Effect of local topographic heterogeneity on tree species assembly in an Acacia-dominated African savanna. Journal of Tropical Ecology 35, 46-56. | 120.00 | 1 |

| | | | |
|---|---|---|---|
| phytosoc. study | Pereira et al. 2003. Use-History Effects on Structure and Flora of Caatinga. Biotropica 35, 154-165. | 0.60 | 1 |
| phytosoc. study | Newton 1988. The structure and phenology of a moist deciduous forest in the Central Indian Highlands. Vegetatio 75, 3-16. | 74.50 | 1 |
| phytosoc. study | Sagar et al. 2003. Tree species composition, dispersion and diversity along a disturbance gradient in a dry tropical forest region of India. Forest Ecology and Management 186, 61-71. | 4.50 | 2 |
| phytosoc. study | Prakasha et al. 2008: Stand Structure of a Tropical Dry Deciduous Forest in Bhadra Wildlife Sanctuary, Karnataka, Southern India. Bulletin of the National Institute of Ecology 19, 1-7. | 2.00 | 1 |
| phytosoc. study | Nguyen & Baker 2016. Structure and composition of deciduous dipterocarp forest in Central Vietnam: patterns of species dominance and regeneration failure. Plant Ecology & Diversity 9 | 2.80 | 1 |
| phytosoc. study | Mohanta et al. 2020. Carbon stock assessment and its relation with tree biodiversity in Tropical Moist Deciduous Forest of Similipal Biosphere Reserve, Odisha, India. Tropical Ecology. https://doi.org/10.1007/s42965-020-00111-8 | 2.10 | 2 |
| phytosoc. study | Sukumar et al. 2004. Mudumalai Forest Dynamics Plot, India. In: Losos & Leigh. Tropical Forest Diversity and Dynamism. The University of Chicago Press, 551-563. | 50.00 | 1 |
| phytosoc. study | Parthasarathy & Karthikeyan 1997. Plant biodiversity inventory and conservation of two tropical dry evergreen forests on the Coromandel coast, south India. Biodiversity and conservation 6, 1063-1083. | 1.00 | 2 |
| phytosoc. study | Deb & Sundriyal 2011. Vegetation dynamics of an old-growth lowland tropical rainforest in North-east India. Species composition and stand heterogeneity. International Journal of Biodiversity and Conservation 3, 405-430. | 4.00 | 1 |
| phytosoc. study | Bannister & Donoso 2013. Forest Typification to Characterize the Structure and Composition of Old-growth Evergreen Forests on Chiloe Island, North Patagonia (Chile). Forests 4, 1087-1105. https://doi.org/10.3390/f4041087 | 4.60 | 1 |
| phytosoc. study | Sadili et al. 2018. Tree species diversity in a pristine montane forest previously untouched by human activities in Foja Mountains, Papua, Indonesia. Reinwardtia 17, 133-154. | 1.00 | 1 |
| phytosoc. study | Sefidi et al. 2015. Effect of topography on tree species composition and volume of coarse woody debris in an Oriental beech (Fagus orientalis Lipsky) old growth forests, northern Iran. iForest 9, 658-665. | 3.50 | 1 |
| phytosoc. study | Rejzek et al. 2016. Loss of a single tree species will lead to an overall decline in plant diversity: Effect of Dracaena cinnabari Balf. f. on the vegetation of Socotra Island. Biological Conservation 196, 165-172. | 6.68 | 1 |
| phytosoc. study | Burns et al. 1999. Dynamics of kahikatea forest remnants in middle North Island: implications for threatened and local plants. Science for Conservation. | 0.16 | 8 |
| phytosoc. study | Linares-Palomino & Ponce-Alvarez 2009. Structural patterns and floristics of a seasonally dry forest in Reserva Ecológica Chaparri, Lambayeque, Peru | 1.00 | 1 |

| | | | |
|---|---|---|---|
| phytosoc. study | Cueva et al. 2019. Efecto de la gradiente altitudinal sobre la composición florística, estructura y biomasa arbórea del bosque seco andino, Loja, Ecuador. Bosque 40, 365-378. | 1.08 | 3 |
| phytosoc. study | Aguirre & Delgado 2005. Vegetación de los bosques secos de Cerro Negro-Cazaderos, Occidente de la Provincia de Loja, 9-24. In: Vásquez et al. Biodiversidad en los Bosques Secos de la Zona de Cerro Negro-Cazaderos, Occidente de la Provincia de Loja: | 1.80 | 1 |
| phytosoc. study | Webb et al. 2011. Structure and Diversity of Seasonal Mixed Evergreen-Deciduous Tropical Forest, Western Thailand. NAT. HIST. BULL. SIAM. SOC. 57 | 1.00 | 1 |
| phytosoc. study | Knab-Vispo et al. 1999. Floristic and structural characterization of a lowland rain forest in the Lower Caura Watershed, Venezuelan Guayana. Acta Botánica Venezuélica 22, 325-359. | 6.70 | 1 |
| phytosoc. study | Teejuntuk et al. 2002. Forest Structure and Tree Species Diversity along an Altitudinal Gradient in Doi Inthanon National Park, Northern Thailand. Tropics 12 | 0.64 | 6 |
| phytosoc. study | García-Villacorta 2009. Diversidad, composición y estructura de un hábitat altamente amenazado: los bosques estacionalmente secos de Tarapoto, Perú. Rev. peru. biol. 16, 081-092. | 0.92 | 1 |
| phytosoc. study | Reddy et al. 2012. Conservation Threat Assessment of Commiphora wightii (Arn.) Bhandari - an Economically Important Species. Taiwania 57, 288-293. | 1.80 | 2 |
| phytosoc. study | Mukherjee 2017. Invasive Prosopis juliflora replacing the Native Floral Community over three decades: a casy study of a World Heritage Site, Keoladeo National Park, India. Biodivers Conserv. 10.1007/s10531-017-1392-y | 26.12 | 1 |
| phytosoc. study | Willems 1999. Forest structure and regeneration dynamics of podocarp/hardwood forest fragments, Banks Peninsula, New Zealand. Master Thesis at Lincoln University. | 2.67 | 1 |
| phytosoc. study | Ahmed & Ogden 1991. Descriptions of some mature Kauri Forests of New Zealand. Tane 33, 89-112. | | 24 |
| phytosoc. study | Kitamura et al. 2005. A botanical inventory of a tropical seasonal forest in Khao Yai National Park, Thailand: implications for fruit-frugivore interactions. Biodiversity and Conservation 14, 1241-1262. | 4.00 | 1 |
| phytosoc. study | Madsen & Ollgaard 1994. Floristic composition, structure, and dynamics of an upper montane rain forest in Southern Ecuador. - Nord. J. Bot. 14: 403-423. Copenhagen. | 1.00 | 2 |
| phytosoc. study | Yumoto et al. 2015. Species composition of a middle altitude forest in Moukalaba-Doudou National Park, Gabon. Tropics 23, 205-213. | 2.25 | 1 |
| phytosoc. study | Lima et al. 2019. Fitossociologia dos componentes lenhose e herbáceo em uma área de caatinga no Cariri Paraibano, PB, Brasil. Hoehnea 46, e792018 | 1.00 | 1 |
| phytosoc. study | Andrade Souza et al. 2019. Phytosociological analysis of the tree-shrub component of the Caatinga, Alagoas, Brazil. Divulgacao Cientifica e Tecnologica Do IFPB, 153-159. | 1.00 | 2 |

| | | | |
|---|---|---|---|
| phytosoc. study | Pinzon et al. 2017. Fine-scale forest variability and biodiversity in the boreal mixedwood forest. Ecography 41, 753-769. | 1.00 | 1 |
| phytosoc. study | Theilade et al. 2022. Evergreen forest types of the central plains in Cambodia: floristic composition and ecological characteristics. Nordic Journal of Botany e03494. https://doi.org/10.1111/njb.03494 | 5.95 | 1 |
| phytosoc. study | Sampaio et al. 2018. Fitossociologia do Cerrado sensu stricto na bacia do Rio Parnaíba no nordeste brasileiro. Advances in Forestry Science 5, 299-307. Location is approximate | 8.00 | 1 |
| phytosoc. study | Felfili et al. 2002. Cmposição florística e fitossociologia do cerrado sentido restrito no município de Água Boa – MT. Acta bot. bras. 16, 103-112. | 1.00 | 1 |
| phytosoc. study | Silva Machado et al. 2019. Florística e fitossociologia de um fragmento de Cerrado lato sensu, Gurupi, TO. Pesquisa Florestal Brasileira. doi: 10.4336/2019.pfb.39e201801685 | 6.70 | 1 |
| phytosoc. study | Sühs et al. 2019. Species diversity, community structure and ecological traits of trees in an upper montane forest, southern Brazil. Acta bot. bras., 153-162. | 1.00 | 1 |
| phytosoc. study | Linares-Palomino et al. 2008. Tree community patterns along a deciduous to evergreen forest gradient in central Bolivia. Ecología en Bolivia, 43(2):1-20. | 1.00 | 3 |
| phytosoc. study | Sadili et al. 2023. Variation in the composition and structure of natural low-land forests at Bodogol, Gunung Gede Pangrango National Park, West Java, Indonesia | 1.50 | 2 |
| phytosoc. study | Simmen et al. 2012. Leaf nutritional quality as a predictor of primate biomass: further evidence of an ecological anomaly within prosimian communities in Madagascar. Journal of Tropical Ecology, 141-151. | 0.37 | 1 |
| phytosoc. study | von Oheimb et al. 2007. Diversity and spatio-temporal dynamics of dead wood in a temperate near-natural beech forest (Fagus sylvatica). Eur J Forest Res, 359-370. | 8.00 | 1 |
| phytosoc. study | Badalamenti et al. 2017. Living and Dead Aboveground Biomass in Mediterranean Forests: Evidence of Old-Growth Traits in a Quercus pubescens Willd. s.l. Stand. Forests, 187. | 1.00 | 1 |
| phytosoc. study | Mori et al. 2007. Roles of disturbance and demographic non-equilibrium in species coexistence, inferred from 25-year dynamics of a late-successional old-growth subalpine forest. Forest Ecology and Management, 74-83. | 1.00 | 1 |
| phytosoc. study | Schwartz et al. 2013. Forest Structure, Stand Composition, and Climate-Growth Response in Montane Forests of Jiuzhaigou National Nature Reserve, China. PLoS One, 371559 | 1.98 | 1 |
| phytosoc. study | Aszalos et al. 2017. First signs of old-growth structure and composition of an oak forest after four decades of abandonment. Biologia, 1264-1274. | 3.00 | 1 |
| phytosoc. study | Ishikawa et al. 1999. Disturbance history and tree establishment in old-growthPinus koraiensis-hardwood forests in the Russian Far East. Journal of Vegetation Science, 439-448. | 0.80 | 2 |
| phytosoc. study | Gilliam & Platt 1999. Effects of long-term fire exclusion on tree species composition and stand structure in an old-growth Pinus palustris (Longleaf pine) forest. Plant Ecology, 15-26. | 21.20 | 2 |

| | | | |
|---|---|---|---|
| phytosoc. study | Motta et al. 2014. Structure, spatio-temporal dynamics and disturbance regime of the mixed beech–silver fir–Norway spruce old-growth forest of Biogradska Gora (Montenegro). Plant Biosystems. https://doi.org/10.1080/11263504.2014.945978 | 1.85 | 1 |
| phytosoc. study | Floyd et al. 2009. Relationship of stand characteristics to drought-induced mortality in three Southwestern pin˜on–juniper woodlands. Ecological Applications, 1223-1230. | 5.30 | 3 |
| phytosoc. study | Gnahore et al. 2023. Floristic composition and structure of closed and open forests in the Banco National Park, Abidjan, Cote d'Ivoire. Asian Journal of Forestry, 17-26. | 4.00 | 1 |
| phytosoc. study | Ahmed et al. 2022. Woody Species Composition, Plant Communities, and Environmental Determinants in Gennemar Dry Afromontane Forest, Southern Ethiopia. Scientifica. Article ID 7970435. https://doi.org/10.1155/2022/7970435 | 1.84 | 1 |
| phytosoc. study | Gwali et al. 2009. Diversity and composition of trees and shrubs in Kasagala forest: a semiarid savannah woodland in central Uganda. African Journal of Ecology, 111-118. | 4.00 | 1 |
| phytosoc. study | Lockhart et al. 2010. Tree Species Composition and Structure in an Old Bottomland Hardwood Forest in South-Central Arkansas. Castanea, 315-329. | 9.84 | 1 |
| phytosoc. study | Ríos, R. C., H. A. Keller & E. R. Krauczuk. 2023. Monodominancia de Calophyllum brasiliense (Calophyllaceae) en sitios de selva higrófila en el NE de Corrientes (Argentina). Bonplandia 32(2): 217-232. Doi: http://dx.doi.org/10.30972/bon.3226437 | 0.80 | 1 |
| phytosoc. study | Riberio et al. 2018. Composition, structure and biodiversity of trees in tropical montane cloud forest patches in Serra do Papagaio State Park, Southeast Brazil. Edinburgh Journal of Botany 75, 255-284. | 2.00 | 1 |
| phytosoc. study | Oliveira et al. 2022. Structure, Biomass and Diversity of a Late-Successional Subtropical Atlantic Forest in Brazil. Floresta e Ambiente 29(4): e20210095 | 3.49 | 1 |
| phytosoc. study | da Rocha et al. 2017. Effect of selective logging on floristic and structural composition in a forest fragment from Amazon Biome. Maringá, v. 39, n. 2, p. 191-199 | 1.25 | 1 |
| phytosoc. study | Pandey et al. 2016. Structure, composition and diversity of forest along the altitudinal gradient in the Himalayas, Nepal. Applied Ecology and Environmental Research 14, 235-251 | 0.20 | 4 |
| phytosoc. study | de Medeiros et al. 2022. Phytosociology of an open arboreal caatinga with high basal area in the Seridó desertification region, Brazil. Rev. Caatinga, Mossoró, v. 36, n. 3, p. 601 – 611, | 0.40 | 1 |
| phytosoc. study | de Lima et al. 2017. Diameter distribution in a Brazilian tropical dry forest domain: predictions for the stand and species. Anais da Academia Brasileira de Ciencias 89, 1189-1203. | 1.60 | 1 |
| phytosoc. study | Kalaba et al. 2013. Floristic composition, species diversity and carbon storage in charcoal and agriculture fallows and management implications in Miombo woodlands of Zambia. Forest Ecology and Management 304, 99-109. | 6.00 | 1 |
| phytosoc. study | Dias Cabacinha et al. 2022. Tree component analysis in a savanna-forest ecotone area of Minas Gerais State. Scientia Agraria Paranaensis, 405-413. | 1.00 | 1 |

| | | | |
|---|---|---|---|
| phytosoc. study | De Souza et al. 2022. Effects of functional traits on the spatial distribution and hyperdominance of tree species in the Cerrado biome. iForest 15, 339-348. 10.3832/ifor3920-015 | 2.16 | 1 |
| phytosoc. study | Scalon et al. 2022. Contrasting strategies of nutrient demand and use between savanna and forest ecosystems in a neotropical transition zone. Biogeosciences 19, 3649-3661. | 2.00 | 1 |
| phytosoc. study | dos Santos, L.O., dos Santos, L.O., de Menezes, M.P.M. et al. Composition and structure of a diverse tree community at the edges of a Brazilian Amazon rainforest island surrounded by marshes and mangroves. Plant Ecol 215, 1469–1481 (2014). https://doi.org/10.1007/s11258-014-0407-y | 1.20 | 1 |
| phytosoc. study | Neto et al. 2022. Structure and tree diversity of an inland atlantic Forest–A case study of ponte branca forest remnant, brazil. The Indonesian Journal of Geography, 54(1), 112-122. doi:https://doi.org/10.22146/ijg.61120 | 2.40 | 1 |
| phytosoc. study | dos Passos et al. 2021. Structure and Tree Diversity in a Mixed Ombrophilous Forest Remnant, Southern Brazil. Floresta e Ambiente 2021; 28(2): e20200064 | 1.00 | 1 |
| phytosoc. study | Pereira et al. 2018. Forest Structure and the Species Composition of the Parque Estadual Mata Atlântica, Located in Goiás State, Brazil. International Journal of Ecology. Article ID 1219374 | | 1 |
| phytosoc. study | He et al. 2015. Forest structure and regeneration of the Tertiary relict Taiwania cryptomerioides in the Gaoligong Mountains, Yunnan, southwestern China. Phytocoenologia Vol. 45 (2015), Issue 1–2, 135–156 | 1.43 | 2 |
| phytosoc. study | Su et al. 2010. Differences in the structure, species composition and diversity of primary and harvested forests on Changbai Mountain, Northeast China. Journal of Forest Science 56, 285-293. | 1.28 | 1 |
| phytosoc. study | Yaguana et al. 2012. Diversidad floristica y estructura del bosque nublado del Rio Numbala, Zamora-Chinchipe, Ecuador: El ‘bosque gigante’ de Podocarpaceae adyacente al Parque Nacional Podocarpus. Revista Amazonica: Ciencia y Tecnologia 1, 226-247. | 1.00 | 1 |
| phytosoc. study | Latt et al. 2022. Tree Species Composition and Forest Community Types along Environmental Gradients in Htamanthi Wildlife Sanctuary, Myanmar: Implications for Action Prioritization in Conservation. Plants 11, 2180. https://doi.org/10.3390/plants11162180 | 4.12 | 1 |
| phytosoc. study | Lusk & Carr 2023. Canopy structure and understorey light availability in Nothofagus and podocarp-broadleaf stands in a New Zealand forest. New Zealand Journal of Botany. DOI: 10.1080/0028825X.2023.2240752 | | 4 |
| phytosoc. study | Merce et al. 2012. The structure of a natural mixed beech – sessile oak forest in Runcu Grosi Natural Reserve. Journal of Horticulture, Forestry and Biotechnology, 131-138. | 3.40 | 1 |
| phytosoc. study | Rugani et al. 2013. Gap Dynamics and Structure of Two Old-Growth Beech Forest Remnants in Slovenia. Plos One. https://doi.org/10.1371/journal.pone.0052641 | 1.57 | 2 |

| | | | |
|---|---|---|---|
| phytosoc. study | Chang et al. 2010. Species Composition, Size-Class Structure, and Diversity of the Lienhuachih Forest Dynamics Plot in a Subtropical Evergreen Broad-Leaved Forest in Central Taiwan. Taiwan J For Sci, 81-95. | 25.00 | 1 |
| phytosoc. study | Chao et al. 2007. Distribution Patterns of Tree Species in the Lanjenchi Lowland Rain Forest. Taiwania, 343-351. | 5.88 | 1 |
| phytosoc. study | Ni et al. 2021. An old-growth subtropical evergreen broadleaved forest suffered more damage from Typhoon Mangkhut than an adjacent secondary forest. Forest Ecology and Management, 119433 | 8.20 | 1 |
| phytosoc. study | Splechtna et al. 2005. Disturbance history of a European old-growth mixed-species forest - A spatial dendro-ecological analysis. Journal of Vegetation Science, 511-522. | 4.00 | 1 |
| phytosoc. study | Vandekerkhove et al. 2018. Very large trees in a lowland old-growth beech (Fagus sylvatica L.) forest: Density, size, growth and spatial patterns in comparison to reference sites in Europe. Forest Ecology and Management, 1-17. | 10.00 | 7 |
| phytosoc. study | Jaloviar et al. 2020. Gap Structure and Regeneration in the Mixed Old-Growth Forests of National Nature Reserve Sitno, Slovakia. Forests 81 | 2.50 | 1 |
| phytosoc. study | Martin-Benito et al. 2022. Development and long-term dynamics of old-growth beech-fir forests in the Pyrenees: Evidence from dendroecology and dynamic vegetation modelling. Forest Ecology and Management, 120541 | 0.99 | 2 |
| phytosoc. study | https://forestgeo.si.edu/sites/europe/zofin | 25.00 | 1 |
| phytosoc. study | Keren et al. 2017. Stand structural complexity of mixed old-growth and adjacent selection forests in the Dinaric Mountains of Bosnia and Herzegovina. Forest Ecology and Management, 531-541. | 1.81 | 2 |
| phytosoc. study | Papadopoulou et al. 2023. Exploring Texture Diversity of Beech-Spruce-Fir Stands through Development Phase Analysis in the Frakto Virgin Forest of Greece. Diversity, 278. | 2.00 | 1 |
| phytosoc. study | Visnjic et al. 2009. Virgin Status Assessment of Plješevica Forest in Bosnia - Herzegovina. Not. Bot. Hort. Agrobot. Cluj, 22-27. | 1.00 | 1 |
| phytosoc. study | Fraver and Palik 2012. Stand and cohort structures of old-growth Pinus resinosa-dominated forests of northern Minnesota, USA. Journal of Vegetation Science, 249-259. | 0.50 | 7 |
| phytosoc. study | Biondi & Bradley 2013. Long-term survivorship of single-needle pinyon (Pinus monophylla) in mixed-conifer ecosystems of the Great Basin, USA. Ecosphere, 120. | 1.00 | 2 |
| phytosoc. study | Barth 2010. Environmental Correlates of Tree Species Distribution in Old-growth Pinus lambertiana—Abies concolor Forests. Bachelor’s Thesis, University of Washington. | 10.20 | 1 |
| phytosoc. study | Lutz et al. 2021. Large-diameter trees, snags, and deadwood in southern Utah, USA. Ecological Processes, 9 | 13.64 | 1 |

| | | | |
|---|---|---|---|
| phytosoc. study | Ansley & Battles 1998. Forest Composition, Structure, and Change in an Old-Growth Mixed Conifer Forest in the Northern Sierra Nevada. The Journal of the Torrey Botanical Society, 297-308. | 4.00 | 1 |
| phytosoc. study | Stephens & Gill 2005. Forest structure and mortality in an old-growth Jeffrey pine-mixed conifer forest in north-western Mexico. Forest Ecology and Management, 15-28. | 4.90 | 1 |
| phytosoc. study | Taylor & Zisheng 1988. Regeneration Patterns in Old-Growth Abies-Betula Forests in the Wolong Natural Reserve, Sichuan, China. Journal of Ecology, 1204-1218. | 3.44 | 1 |
| phytosoc. study | Taylor et al. 2005. Regeneration patterns and tree species coexistence in old-growth Abies–Picea forests in southwestern China. Forest Ecology and Management, 303-317 | 1.90 | 1 |
| phytosoc. study | Rhoades et al. 2017. A Decade of Streamwater Nitrogen and Forest Dynamics after a Mountain Pine Beetle Outbreak at the Fraser Experimental Forest, Colorado. Ecoystems, 380–392. https://doi.org/10.1007 | 2.14 | 1 |
| phytosoc. study | Dai et al. 2011. Changes in forest structure and composition on Changbai Mountain in Northeast China. Annals of Forest Science, 889-897. | 1.00 | 2 |
| phytosoc. study | Sharma et al. 2017. Effect of altitudinal gradients on forest structure and composition on ridge tops in Garhwal Himalya. Energ. Ecol. Environ., 404-417. DOI 10.1007/s40974-017-0067-6 | 0.60 | 5 |
| phytosoc. study | Parveen & Ilyas 2022: Tree diversity and natural regeneration in Tropical Dry Deciduous Forest of Panna Tiger Reserve, India. Preprint on researchsquare.com. https://doi.org/10.21203/rs.3.rs-629351/v1 | 10.61 | 1 |
| phytosoc. study | Malimbwi et al. 1994: Estimation of Biomass and Volume in Miombo Woodland at Kitulangalo Forest Reserve, Tanzania. Journal of Tropical Forest Science 7, 230-242. | | 1 |
| phytosoc. study | Joshi et al. 2021. Tree biomass and carbon stock assessment of subtropical and temperate forests in the Central Himalaya, India. Trees, Forests and People 6, 100147 | 0.50 | 4 |
| phytosoc. study | Temesgen & Warkineh 2020. Woody Species Structure and Regeneration Status in Kafta Sheraro National Park Dry Forest, Tigray Region, Ethiopia. International Journal of Forestry Research, 22 pages. | 6.44 | 1 |
| phytosoc. study | Das 2024. Tree species diversity, composition and structure in the tropical moist deciduous forest of Kadigarh National Park, Mymensingh, Bangladesh. Asian Journal of Forestry 8, 41-49 | 2.48 | 1 |
| phytosoc. study | Ouoba & Constant 2019. Woody plants structure and composition in Burkina Faso Sahel: case study in Kékénéné village. Int. J. Biol. Chem. Sci. 13, 1682-1692. | 31.00 | 1 |
| phytosoc. study | Muledi et al. 2017. Fine-scale habitats influence tree species assemblage in a miombo forest. Journal of Plant Ecology 10, 958–969. | 10.00 | 1 |
| phytosoc. study | Michela & Galindez 2021. Characterization of a forest in the center west of the province of Chaco, Argentina. Sustainable Forestry 4. doi:10.24294/sf.v4i2.1611 | 1.00 | 1 |

| | | | |
|---|---|---|---|
| phytosoc. study | Alem & Woldemariam 2009. A comparative assessment on regeneration status of indigenous woody plants in Eucalyptus grandis plantation and adjacent natural forest. Journal of Forestry Research 20, 31-36. | 0.80 | 1 |
| phytosoc. study | Parker et al. 1984. Tree dynamics in an old-growth, deciduous forest. Forest Ecology and Management, 31-57. | 20.60 | 1 |
| phytosoc. study | Tavankar 2015. Structure of natural Juniperus excelsa stands in Northwest of Iran. Biodiversitas, 161-167. | 2.20 | 1 |
| phytosoc. study | Nouri et al. 2015. Comparison of woody species diveristy between managed and unmanaged forests considering vertical structure in Hyrcanian forests, Iran. Biodiversitas 16, 95-101. | 4.00 | 1 |
| phytosoc. study | Amiri & Naghdi 2016. Assessment of competition indices of an unlogged oriental beech mixed stand in Hyrcanian forests, Northern Iran. Biodiversitas 17, 306-314. | 16.00 | 1 |
| phytosoc. study | Adel et al. 2013. Forest structure and woody plant species composition after a wildfire in beech forests in the north of Iran. Journal of Forestry Research 24, 255-262. | 6.00 | 1 |
| phytosoc. study | Huan-Yu Lin et al. 2005. Species Composition and Structure of a Montane Rainforest of Mt. Lopei in Northern Taiwan. Taiwania, 234-249. | 1.00 | 1 |
| phytosoc. study | Chai et al. 2017. Population structure and spatial pattern of predominant tree species in a pine–oak mosaic mixed forest in the Qinling Mountains, China. Journal of Plant Interactions. | 1.20 | 1 |
| phytosoc. study | Tang et al. 2020. Forest characteristics, population structure and growth trends of Pinus yunnanensis in Tianchi National Nature Reserve of Yunnan, southwestern China. Vegetation Classification and Survey 1: 7-20. https://doi.org/10.3897/VCS/2020/37980 | 1.29 | 1 |
| phytosoc. study | Sulistyawati et al. 2018. Tree Community Structure and Composition of a One-Hectare Permanent Plot in the Montane Zone of Mount Kerinci, Kerinci Seblat National Park, Jambi | 1.00 | 1 |
| phytosoc. study | Khamyong, S., Lykke, A.M., Seramethakun, D. and Barfod, A.S. (2003), Species composition and vegetation structure of an upper montane forest at the summit of Mt. Doi Inthanon, Thailand. Nordic Journal of Botany, 23: 83-97. https://doi.org/10.1111/j.1756-1051.2003.tb00371.x | 8.00 | 1 |
| phytosoc. study | Hamann et al. 1999. A botanical inventory of a submontane tropical rainforest on Negros Island, Philippines. Biodiversity and Conservation 8, 1017-1031 | 1.00 | 1 |
| phytosoc. study | Gilbert et al. 2010. Beyond the tropics: forest structure in a temperate forest mapped plot. Journal of Vegetation Science, 388-405. | 6.00 | 1 |
| phytosoc. study | Dagley 2008. Spatial pattern of coast redwood in three alluvial flat old-growth forests in Northern California. Forest Science 54 (3) | 1.50 | 3 |
| phytosoc. study | Ao et al. 2023. Stand Structure, Regeneration Potential and Biomass Carbon Stock of Subtropical Forest of Mizoram, Northeast India. Indian Journal of Ecology 50, 1972-1979. | 0.40 | 1 |

| | | | |
|---|---|---|---|
| phytosoc. study | Marziliano et al. 2017. Forest structure of a maple old-growth stand: a case study on the Apennines mountains (Southern Italy). Journal of Mountain Science 14. DOI: 10.1007/s11629-016-4336-1 | 0.50 | 1 |
| phytosoc. study | Takahashi et al. 2007. Quantitative and qualitative effects of a severe ice storm on an old-growth beech–maple forest. Canadian Journal of Forest Research, 598-606. https://doi.org/10.1139/X06-266 | 0.98 | 1 |
| phytosoc. study | Orwig et al. 2022. Land-use history impacts spatial patterns and composition of woody plant species across a 35-hectare temperate forest plot. PeerJ 10:e12693 https://doi.org/10.7717/peerj0.12693 | 35.00 | 1 |
| phytosoc. study | Brzeziecki et al. 2018. Structural and compositional dynamics of strictly protected woodland communities with silvicultural implications, using Białowieża Forest as an example. Annals of Forest Science, 89 | 15.44 | 1 |
| phytosoc. study | Ruschel et al. 2006. Woody plant species richness in the Turvo State park, a large remnant of deciduous Atlantic forest, Brazil | | 1 |
| phytosoc. study | Abe 2007. Forest management impacts on growth, diversity and nutrient cycling of lowland tropical rainforest and plantations, Papua New Guinea. Doctoral Thesis, The University of Western Australia, School of Plant Biology. | 2.00 | 1 |
| phytosoc. study | Ball et al. 2019. Data from a botanical survey of the Anogeissus cloud forest in Jabal Qamar, Dhofar, Oman. Mendeley Data, V1, doi: 10.17632/dc97zn6gzc0.1 | | 30 |
| phytosoc. study | Borah et al. 2015. Tree species composition, biomass and carbon stocks in two tropical forest of Assam. Biomass and Bioenergy 78, 25-35. | | 2 |
| phytosoc. study | Marod et al. 2022. Population Structure and Spatial Distribution of Tree Species in Lower Montane Forest, Doi Suthep-Pui National Park, Northern Thailand. Environment and Natural Resources Journal, 644-663. | 16.00 | 1 |
| phytosoc. study | Chandrashekara & Ramakrishnan 1994. Vegetation and gap dynamics of a tropical wet evergreen forest in the Western Ghats of Kerala, India. Journal of Tropical Ecology 10, 337-354. | 5.00 | 1 |
| phytosoc. study | Bordin et al. 2019. Community structure and tree diversity in a subtropical forest in southern Brazil. Biota Neotropica. 19(2): e20180606. http://dx.doi.org/10.1590/1676-0611-BN-2018-0606 | 1.20 | 1 |
| phytosoc. study | de Melo 2013. Relationships between structure of the tree component and environmental variables in a subtropical seasonal forest in the upper Uruguay River valley, Brazil. Acta Amazonica Brasicila 27, 751-760. | 1.00 | 1 |
| phytosoc. study | Simmen et al. 2005. Richesse en métabolites secondaires des forêts de Mayotte et de Madagascar et incidence sur la consommation de feuillage chez deux espèces de lémurs. Revue d’Ecologie, Terre de Vie, 297-324. | 0.90 | 5 |

| | | | |
|---|---|---|---|
| phytosoc. study | Paiva et al. 2021. Fitossociologia da caatinga na Floresta Nacional de Açu, Estado do Rio Grande do Norte, Brasil, e entorno: diversidade e biogeografia do componente lenhoso. Hoehnea 48: e222020. https://doi.org/10.1590/2236-8906-22/2020 | | 1 |
| phytosoc. study | Kittur et al. 2014. Wildland fires and moist deciduous forests of Chhattisgarh, India: divergent component assessment. Journal of Forestry Research 25, 857-866. | 3.20 | 1 |
| phytosoc. study | Kumar et al. 2011. Forest structure, diversity and soil properties in a dry tropical forest in Rajasthan, Western India. Annals of Forest Research 54, 89-98. | 1.00 | 1 |
| phytosoc. study | Naidu et al. 2018. Assessment of tree diversity in tropical deciduous forests of Northcentral Eastern Ghats, India, Geology, Ecology, and Landscapes 2, 216-227. DOI: 10.1080/24749508.2018.1452479 | 1.00 | 6 |
| phytosoc. study | Su et al. 2007. Fushan subtropical forest dynamics plot: tree species characteristics and distribution patterns. Taipei: Taiwan Forestry Research Institute. | 25.00 | 1 |
| phytosoc. study | Carvalho, D. A. et al. 2005. Variações florísticas e estruturais do componente arbóreo de uma floresta ombrófila alto-montana às margens do rio Grande, Bocaina de Minas, MG, Brasil. Acta Bot. Brasil. 19(1): 91–109 | 1.20 | 1 |
| phytosoc. study | Pompeu et al. 2014. Floristic composition and structure of an upper montane cloud forest in the Serra da Mantiqueira Mountain Range of Brazil. Acta Botanica Brasilica 28(3): 456-464. 2014 | 0.60 | 1 |
| phytosoc. study | Assunçao & Felfili 2004. Fitossociologia de um fragmento de cerrado sensu stricto na APA do Paranoá, DF, Brasil. Acta bot. bras. 18, 903-909. | 1.00 | 1 |
| phytosoc. study | Rossi et al. 1998. Fitossociologia do estrato arbóreo do cerrado (sensu stricto) no parque ecológiqo norte, Brasília - DF. Bol. Herb. Ezechias Paulo Heringer 2, 49-56. | 1.00 | 1 |
| phytosoc. study | Armesto & Figueroa 1987. Stand Structure and Dynamics in the Temperate Rain Forests of Chiloe Archipelago, Chile. Journal of Biogeography 14, 367-376. | 0.32 | 1 |
| phytosoc. study | Mensah et al. 2018. Vegetation structure, dominance patterns and height growth in an Afromontane forest, Southern Africa. J. For. Res. https://doi.org/10.1007/s11676-018-0801-8 | 1.50 | 1 |
| phytosoc. study | Igu 2023. Species Distribution and Patterns in a Forest-savannah Ecotone: Environmental Change and Conservation Concerns. Journal of Botanical research 5, 27-35. https://doi.org/10.30564/jbr.v5i3.5588 | 2.00 | 1 |
| phytosoc. study | Kanagaraj 2017. Assessment of tree species diversity and its distribution pattern in Pachamalai Reserve Forest, Tamil Nadu. Journal of Sustainable Forestry 36, 32-46, DOI: 10.1080/10549811.2016.1238768 | 0.45 | 1 |
| phytosoc. study | Guimarães Giácomo et al. 2013. Florística e fitossociologia em áreas de campo sujo e cerrado sensu stricto na estação ecológica de pirapitinga - MG. Ciência Florestal, Santa Maria 23, 29-43. | 1.90 | 1 |

| | | | |
|---|---|---|---|
| phytosoc. study | Manikandan et al. 2019. Wood Stem Density and Above-ground Biomass in Pachaimalai Hills of Southern Estern Ghats, Tamil Nadu, India. International Journal for Research in Applied Science & Engineering Technology 7 | 10.00 | 1 |
| phytosoc. study | Kozera et al. 2005. Fitossociologica do componente arbóreo de um fragmento de floresta ombrófila mista montana, Curitiba, PR, BR1. Floresta 36. | | 1 |
| phytosoc. study | Pansonato et al. 2019. Community structure and species composition of a periodically flooded Restinga forest in Caraguatatuba, São Paulo, Brazil. Biota Neotropica. 19(1): e20170477. http://dx.doi.org/10.1590/1676-0611-BN-2017-0477 | 1.60 | 1 |
| phytosoc. study | Semegnew et al. 2021. Woody Species Composition, Vegetation Structure, and Regeneration Status of Majang Forest Biosphere Reserves in Southwestern Ethiopia. International Journal of Forestry Research. Article ID 5534930, 22 pages. https://doi.org/10.1155/2021/5534930 | 1.40 | 4 |
| phytosoc. study | Evitex-Izayas & Udayakumar 2021. Density, diversity and community composition of trees in tropical thorn forest, peninsular India. Current Botany 12, 138-145. | 1.00 | 1 |
| phytosoc. study | Nagaraj & Udayakumar 2021. Aboveground Biomas Stockpile of Trees in Southern Thorn Forest, Tuticorin, Peninsular India. Current World Environment 16, 755-763. | 1.00 | 1 |
| phytosoc. study | Godlee et al. 2020: Diversity and Structure of an Arid Woodland in Southwest Angola, with Comparison to the Wider Miombo Ecoregion. Diversity 12, 140. BA has been backadjusted to full plot area. | 15.00 | 1 |
| phytosoc. study | Gonçalves et al. 2018. Tree Species Diversity and Composition of Miombo Woodlands in South-Central Angola: A Chronosequence of Forest Recovery after Shifting Cultivation. International Journal of Forestry Research, Article ID 6202093. https://doi.org/10.1155/2017/6202093 | 1.00 | 1 |
| phytosoc. study | Oyewolo 2023. Phytosociological Assessment and Diversity of Woody Species in Omo Biosphere Reserve, Nigeria. Journal of the Cameroon Academy of Sciences 19, 125-139. | 3.00 | 1 |
| phytosoc. study | Githae et al. 2007. A botanical inventory and diversity assessment of Mt Marsabit forest, a sub-humid montane forest in the arid lands of northern Kenya. African Journal of Ecology 46, 39-45. | 1.50 | 1 |
| phytosoc. study | Yadav & Gupta 2006. Effect of micro-environment and human disturbance on the diversity of woody species in the Sariska Tiger Project in India. Forest Ecology and Management 225, 178-189. | 1.30 | 1 |
| phytosoc. study | Kakkar et al. 2021. Patterns of woody species diversity and structure in Thalewood House permanent preservation plot in Bannerghatta National Park, Bangalore, India. Tropical Ecology. https://doi.org/10.1007/s42965-021-00169-y | 1.00 | 1 |
| phytosoc. study | Workayehu et al. 2022. Floristic Composition, Diversity, and Vegetation Structure of Woody Species in Kahitassa Forest, Northwestern Ethiopia. International Journal of Forestry Research. Article ID 7653465. https://doi.org/10.1155/2022/7653465 | 4.04 | 1 |

| | | | |
|---|---|---|---|
| phytosoc. study | Young et al. 2017. Variation in population structure and dynamics of montane forest tree species in Ethiopia. Guide priorities for conservation and research. Biotropica 49, 309-317. https://doi.org/10.1111/btp.12420 | | 4 |
| phytosoc. study | Duarte Barbosa et al. 2012. Florística e fitossociologia de espécies arbóreas e arbustivas em uma área de caatinga em Arcoverde, PE, Brasil. Revista árvore 36, 851-858. | 1.00 | 1 |
| phytosoc. study | Pourbabaei et al. 2014. Comparison in woody species composition, diversity and community structure as affected by livestock grazing and human uses in beech forests of Northern Iran. Forestry Ideas 20. | 2.50 | 1 |
| phytosoc. study | Wiseman et al. 2004. Woody vegetation change in response to browsing in Ithala Game Reserve, South Africa. South African Journal of Wildlife Research 34, 25-37. | | 1 |
| phytosoc. study | Saiter et al. 2011. Tree changes in a mature rainforest with high diversity and endemism on the Brazilian coast. Biodivers Conserv 20,1921–1949 | 1.02 | 1 |
| phytosoc. study | Mereiles Monteiro & Coutinho Kurtz 2020. Phytosociology of Two Caatinga Phytophysiognomies with Different Histories of Anthropic Disturbance. Floresta Ambient. 27 | 2.40 | 1 |
| phytosoc. study | Donoso & Soto 2016. Does site quality affect the additive basal area phenomenon? Results from Chilean old-growth temperate rainforests. Can. J. For. Res. 46, 1330-1336 | 6.10 | 2 |
| phytosoc. study | Gutiérrez et al. 2004. Disturbance and regeneration dynamics of an old-growth North Patagonian rain forest in Chiloé Island, Chile. Journal of Ecology 92: 598-608. https://doi.org/10.1111/j.0022-0477.2004.00891.x | 0.20 | 1 |
| phytosoc. study | Josse C. & Balslev H. 1994. The composition and structure of a dry, semideciduous forest in western Ecuador. - Nord. J. Bot. 14: 425434 | 1.00 | 1 |
| phytosoc. study | Ribeiro da Costa & Soares de Araújo 2007. Organização comunitária de um encrave de cerrado sensu stricto no bioma Caatinga, chapada do Araripe, Barbalha, Ceará. Acta bot. bras 21, 281-291. | | 1 |
| phytosoc. study | Da Cruz Silva et al. 2016. Florística, fitossociologia e caracterização sucessional em um remanescente de Caatinga em Sergipe. Gaia Scientia 10, 1-14. | 1.20 | 1 |
| phytosoc. study | Souza Miranda et al. 2002. Community Structure of Woody Plants of Roraima Savannahs, Brazil. Plant Ecology 164, 109-123. | 6.75 | 1 |
| phytosoc. study | Tadros & Ananbeh 2018. Vegetation Composition and Structure of Woody Plant Communities in Ajloun Forest Reserve. Jordan Journal of Agricultural Sciences. | | 1 |
| phytosoc. study | Triepke et al. 2012. Composition and Structure of Aleppo Pine (Pinus halepensis) Communities in the Dibeen Forest Reserve, Jordan, 356-366 https://doi.org/10.3375/043.032.0403. | 1.45 | 1 |
| phytosoc. study | López Serrano et al. 2022. Diversity and ecological importance of tree vegetation at El Tecuan Park in the state of Durango, Mexico. Revista Mexicana de ciencias forestales. https://doi.org/10.29298/rmcf.v13i74.1273 | 16.80 | 1 |

| | | | |
|---|---|---|---|
| phytosoc. study | Davidar et al. 2007. Floristic inventory of woody plants in a tropical montane (shola) forest in the Palni hills of the Western Ghats, India. Tropical Ecology 48, 15-25. | 1.08 | 1 |
| phytosoc. study | Worku et al. 2022. Diversity, Structural, and Regeneration Analysis of Woody Species in the Afromontane Dry Forest of Harego, Northeastern Ethiopia. International Journal of Forestry Research. Article ID 7475999. Https://doi.org/10.1155/2022/7475999 | 2.88 | 1 |
| phytosoc. study | Bonino & Araujo 2005. Structural differences between a primary and a secondary forest in the Argentine Dry Chaco and management implications. Forest Ecology and Management 206, 407-412. | 1.60 | 1 |
| phytosoc. study | Meireles & Shepherd 2015. Structure and floristic similarities of upper montane forests in Serra Fina mountain range, southeastern Brazil. Acta bot. bras. 29, 58-72. | 0.30 | 1 |
| phytosoc. study | Gonçalves et al. 2018b. Species diversity, population structure and regeneration of woody species in fallows and mature stands of tropical woodlands of southeast Angola. J. For. Res. 29, 1569–1579. https://doi.org/10.1007/s11676-018-0593-x | 1.00 | 1 |
| phytosoc. study | Yadav 2016. Species structure and diversity in Achanakmar-Amarkantak Biosphere reserve, Central India. Journal of Applied and Natural Science 8, 1241 - 1248 | 1.00 | 1 |
| phytosoc. study | Rawat et al. 2010. Diversity, distribution and vegetation assessment in the Jahlmanal watershed in cold desert of the Lahaul valley, north-western Himalaya, India. iForest 3, 65-71. | 1.00 | 1 |
| phytosoc. study | Baltzer et al. 2021. Permafrost thaw in boreal peatlands is rapidly altering forest community composition. Dryad Dataset, https://doi.org/10.5061/dryad.0cfxpnw0p | 9.60 | 1 |
| phytosoc. study | Siitonen et al. 2000. Coarse woody debris and stand characteristics in mature managed and old-growth boreal mesic forests in southern Finland. Forest Ecology and Management, 211-225. | 9.00 | 1 |
| phytosoc. study | Kuuluvainen et al. 2002. Tree age distributions in old-growth forest sites in Vienansalo wilderness, eastern Fennoscandia. Silva Fennica, 169–184. | 1.60 | 1 |
| phytosoc. study | Eichhorn 2010. Boreal Forests of Kamchatka: Structure and Composition. Forests, 154-176. doi:10.3390/f1030154 | 0.25 | 8 |
| phytosoc. study | Yu et al. 2009. The impact of fire and density-dependent mortality on the spatial patterns of a pine forest in the Hulun Buir sandland, Inner Mongolia, China. Forest Ecology and Management, 2098-2107. | 1.00 | 1 |
| phytosoc. study | Pouta et al. 2022. Partitioning of Space Among Trees in an Old-Growth Spruce Forest in Subarctic Fennoscandia. Front. For. Glob. Change, 09 June 2022. Sec. Forest Management. Volume 5 - 2022 \| https://doi.org/10.3389/ffgc.2022.817248 | 8.80 | 1 |
| phytosoc. study | Khakimulina et al. 2015. Mixed-severity natural disturbance regime dominates in an old-growth Norway spruce forest of northwest Russia. Journal of Vegetation Science, 400-413. | 1.80 | 1 |

| | | | |
|---|---|---|---|
| phytosoc. study | Ding et al. 2019. Intraspecific trait variation and neighborhood competition drive community dynamics in an old-growth spruce forest in northwest China. Science of the Total Environment, 525-532. | 15.00 | 1 |
| phytosoc. study | Mwampashi 2013. Wood land structure, basic density and above ground carbon stock estimations of wet miombo woodlands in Mbozi district Tanzania. Master thesis. | 2.96 | 1 |
| phytosoc. study | Kafle 2004. Effects of Forest Fire Protection on Plant Diversity in a Tropical Deciduous Dipterocarp-Oak Forest, Thailand. Proceedings of the Second International Symposium on Fire Economics, Planning, and Policy: A Global View | 4.00 | 1 |
| phytosoc. study | Oakes et al. 2014. Long-term vegetation changes in a temperate forest impacted by climate change. Ecosphere, 135. http://dx.doi.org/10.1890/ES14-00225.1 | 0.67 | 1 |
| phytosoc. study | Sitati et al. 2016. Tree Species Diversity and Dominance in Ketumbeine Forest Reserve, Tanzania. Journal of Biodiversity Management and Forestry 5, 3 | 5.47 | 1 |
| phytosoc. study | Pala et al. 2011. Species composition and phytosociological status of Chanderbadni Sacred forest in Garwhal Himalaya, Uttarakhand India. NeBIO 2, 52-59. | 1.00 | 1 |
| phytosoc. study | Yineger et al. 2008. Floristic composition and structure of the dry afromontane forest at Bale Mountains National Park, Ethiopia. Ethiop. J. Sci 31, 103-120. | 6.12 | 1 |
| phytosoc. study | Martin-Benito et al. 2020. Disturbances and Climate Drive Structure, Stability, and Growth in Mixed Temperate Old-growth Rainforests in the Caucasus. Ecosystems, 1170-1185. | 1.41 | 1 |
| phytosoc. study | Bolte et al. 2013. Space sequestration below ground in old-growth spruce-beech forests—signs for facilitation? Front Plant Sci, 322 | 1.00 | 3 |
| phytosoc. study | Hussain et al. 2010. Phytosociology and Structure of Central Karakoram National Park (CKNP) of Northern Areas of Pakistan. World Applied Sciences Journal, 1443-1449. | | 5 |
| phytosoc. study | Sione, S.M.J., Wilson, M.G., Ledesma, S.G. et al. Driving factors of tree biomass and soil carbon pool in xerophytic forests of northeastern Argentina. Ecol Process 12, 64 (2023). https://doi.org/10.1186/s13717-023-00478-1 | 1.80 | 1 |
| phytosoc. study | Fajardo & de Graaf 2004. Tree dynamics in canopy gaps in old-growth forests of Nothofagus pumilio in Southern Chile. Plant Ecology 173, 95-105. | | 2 |
| phytosoc. study | Keyes and Teraoka 2014. Structure and Composition of Old-Growth and Unmanaged Second-Growth Riparian Forests at Redwood National Park, USA. Forests, 5, 256-268; doi:10.3390/f5020256 | 0.16 | 1 |
| phytosoc. study | Pouta et al. 2022. Partitioning of Space Among Trees in an Old-Growth Spruce Forest in Subarctic Fennoscandia. Front. For. Glob. Change, 09 June 2022. Sec. Forest Management. Volume 5 - 2022 \| https://doi.org/10.3389/ffgc.2022.817248 | 8.80 | 1 |
| phytosoc. study | Popa et al. 2017. Stand structure, recruitment and growth dynamics in mixed subalpine spruce and Swiss stone pine forests in the Eastern Carpathians. Science of the Total Environment, 1050-1057. | 3.59 | 1 |

| | | | |
|---|---|---|---|
| phytosoc. study | Tiscar & Luca-Borja 2016. Structure of old-growth and managed stands and growth of old trees in a Mediterranean Pinus nigra forest in southern Spain. Forestry, 201-207. | 1.06 | 1 |
| phytosoc. study | Taylor 2010. Fire disturbance and forest structure in an old-growth Pinus ponderosa forest, southern Cascades, USA. Journal of Vegetation Science, 561-572. | 3.46 | 2 |
| field inventory | NEON (National Ecological Observatory Network). Vegetation structure (DP1.10098.001), RELEASE-2023. https://doi.org/10.48443/73zn-k414. Dataset accessed from https://data.neonscience.org on March 10, 2023 | 0.04 | 412 |
| field inventory | Urrutia-Jalabert et al. 2016: Data from: The oldest, slowest forests in the world? Exceptional biomass and slow carbon dynamics of Fitzroya cupressoides temperate rainforests in southern Chile. Dryad Dataset. https://doi.org/10.5061/dryad.2kh91 | 1.20 | 2 |
| field inventory | Kohyama et al. 2020. Trade-off between standing biomass and productivity in species-rich tropical forest: Evidence, explanations and implications. J Ecol. 108, 2571– 2583. https://doi.org/10.1111/1365-2745.13485, https://github.com/kohyamat/p-B. Species codes from Kohyama et al. 2014. | 50.00 | 1 |
| field inventory | Schlund et al. 2015. TanDEM-X data for aboveground biomass retrieval in a tropical peat swamp forest. Remote Sensing of Environment, 158, 255–266. Extracted from Tallo database, Jucker et al. 2022, in preparation. | 0.80 | 1 |
| field inventory | Gentry transects. Data compiled by Alwyn H. Gentry and Missouri Botanical Garden. https://www.mobot.org/MOBOT/Research/gentry/transect.shtml | 0.10 | 195 |
| field inventory | Humboldt Institute, Colombia 2013, collectionCode: IAvH, recorded by: Roy Gonzalez-M.; Jhon Nieto; Daniel García | 1.00 | 4 |
| field inventory | Joint Remote Sensing Research Program (2016): Biomass Plot Library - National collation of stem inventory data and biomass estimation, Australian field sites. Version 1.0.0. Terrestrial Ecosystem Research Network (TERN). Original source: CSIRO | 0.18 | 180 |
| field inventory | Joint Remote Sensing Research Program (2016): Biomass Plot Library - National collation of stem inventory data and biomass estimation, Australian field sites. Version 1.0.0. Terrestrial Ecosystem Research Network (TERN). Original source: DELWP Victoria | 0.04 | 266 |
| field inventory | Joint Remote Sensing Research Program (2016): Biomass Plot Library - National collation of stem inventory data and biomass estimation, Australian field sites. Version 1.0.0. Terrestrial Ecosystem Research Network (TERN). Original source: Department of the Environment Australia | 0.09 | 27 |
| field inventory | Joint Remote Sensing Research Program (2016): Biomass Plot Library - National collation of stem inventory data and biomass estimation, Australian field sites. Version 1.0.0. Terrestrial Ecosystem Research Network (TERN). Original source: DSITI Queensland Herbarium | 0.32 | 950 |
| field inventory | Joint Remote Sensing Research Program (2016): Biomass Plot Library - National collation of stem inventory data and biomass estimation, Australian field sites. Version 1.0.0. Terrestrial Ecosystem Research Network (TERN). Original source: DSITI Queensland RSC | 0.25 | 20 |

| | | | |
|---|---|---|---|
| field inventory | Joint Remote Sensing Research Program (2016): Biomass Plot Library - National collation of stem inventory data and biomass estimation, Australian field sites. Version 1.0.0. Terrestrial Ecosystem Research Network (TERN). Original source: TERN Australia | 1.00 | 53 |
| field inventory | Joint Remote Sensing Research Program (2016): Biomass Plot Library - National collation of stem inventory data and biomass estimation, Australian field sites. Version 1.0.0. Terrestrial Ecosystem Research Network (TERN). Original source: UQ Joint Remote Sensing Research Program | 0.50 | 18 |
| field inventory | Feeley et al. 2013: Compositional shifts in Costa Rican forests due to climate-driven species migrations. Global Change Biology 2013, 3472-3480. | 1.00 | 10 |
| field inventory | Cook et al. 2011: NACP New England and Sierra National Forests Biophysical Measurements: 2008-2010. ORNL DAAC Oak Ridge, Tennessee, USA. | 1.00 | 44 |
| field inventory | Franklin et al. 2015. Data from: Regional variation in Caribbean dry forest tree species composition, Dryad, Dataset, https://doi.org/10.5061/dryad.2r5r9 | 0.06 | 25 |
| field inventory | Potts et al. 2017: Data from: Habitat patterns in tropical rain forests: a comparison of 105 plots in northwest Borneo. Dryad Dataset. https://doi.org/10.5061/dryad.64d74. But refers back to old Potts et al. 2002 publication. | 13.20 | 14 |
| field inventory | Osuri et al. 2019: Data from: Effects of restoration on tree communities and carbon storage in rainforest fragments of the Western Ghats, India. Dryad Dataset. https://doi.org/10.5061/dryad.g7j45sn | 0.08 | 7 |
| field inventory | Condit et al. 2013. Data from Tree Censuses and Inventories in Panama. Smithsonian website. | 1.00 | 36 |
| field inventory | Ferry et al. 2021. Long-term high densities of African elephants clear the understorey and promote a new stable savanna woodland community. https://doi.org/10.5281/zenodo.5564836 | 0.25 | 6 |
| field inventory | Phillips et al. 2003.Efficient plot-based floristic assessment of tropical forests. Journal of Tropical Ecology 19, 629-645. doi:10.1017/S0266467403006035. | 0.30 | 24 |
| field inventory | Goncalves et al. 2018. Tree Inventory and Biometry Measurements, Tapajos National Forest, Para, Brazil, 2010. ORNL DAAC, Oak Ridge, Tennessee, USA. https://doi.org/10.3334/ORNLDAAC/1552. | 0.50 | 4 |
| field inventory | Clark et al. 2016. CARBONO project, La Selva. https://tropicalstudies.org/carbono-project/#1554994314095-2707918b-e855 | 9.00 | 1 |
| field inventory | Craven et al. 2018. OpenNahele: the open Hawaiian forest plot database. Biodiversity Data Journal 6: e28406. https://doi.org/10.3897/BDJ.6.e28406 | 0.07 | 377 |
| field inventory | Ramesh et al. 2010. Forest stand structure and composition in 96 sites along environmental gradients in the central Western Ghats of India. Ecology, 91: 3118-3118. doi:10.1890/10-0133.1 | 1.00 | 32 |

| | | | |
|---|---|---|---|
| field inventory | Miesner et al. 2022. Tree data set from forest inventories in north-eastern Siberia. PANGAEA, https://doi.org/10.1594/PANGAEA.943547 | 0.07 | 122 |
| field inventory | Kohyama et al. 2023. Contribution of tree community structure to forest productivity across a thermal gradient in eastern Asia. Nat. Commun. 14, 1113. https://doi.org/10.1038/s41467-023-36671-1 & https://zenodo.org/records/7668416 | 1.00 | 52 |
| field inventory | McMahon et al. 2023. SERC ForestGEO inventory census data. Smithsonian Environmental Research Center. Dataset. https://doi.org/10.25573/serc.24747231.v2 | 16.00 | 1 |
| field inventory | Pennington et al. 2019. Forest plot inventory data from seasonally dry and moist Atlantic forest in Rio de Janeiro State, 2015-2017 NERC Environmental Information Data Centre. https://doi.org/10.5285/aa3babe9-072c-42ce-9ea5-9dbb921a922d | | 32 |
| field inventory | Franklin et al. 2023. Long-term growth, mortality and regeneration of trees in permanent vegetation plots in the Pacific Northwest, 1910 to present ver 24. Environmental Data Initiative. https://doi.org/10.6073/pasta/5835a1fadb7fd65842f90256edae999c (Accessed 2024-04-03). | 1.00 | 102 |
| field inventory | Maleki, Kobra et al. 2020. A 249-year chronosequence of forest plots from eight successive fires in the eastern Canada boreal mixedwoods Dryad Dataset, https://doi.org/10.5061/dryad.tqjq2bvwz. | 1.00 | 5 |
| field inventory | Hollinger et al. 2021. Howland Forest 25-year dataset - Multi-decadal carbon cycle measurements indicate resistance to external drivers of change at the Howland Forest AmeriFlux site. Fort Collins, CO: Forest Service Research Data Archive. https://doi.org/10.2737/RDS-2021-0014 | 3.00 | 1 |
| field inventory | Wang et al. 2017. Individual size variation reduces spatial variation in abundance of tree community assemblage, not of tree populations. Ecol Evol. 7, 10815– 10828. https://doi.org/10.1002/ece3.3594 | 0.25 | 13 |
| field inventory | Joint Remote Sensing Research Program (2016): Biomass Plot Library - National collation of stem inventory data and biomass estimation, Australian field sites. Version 1.0.0. Terrestrial Ecosystem Research Network (TERN). Original source: Western Regeneration Pty. Ltd. | 0.10 | 73 |
| field inventory | Joint Remote Sensing Research Program (2016): Biomass Plot Library - National collation of stem inventory data and biomass estimation, Australian field sites. Version 1.0.0. Terrestrial Ecosystem Research Network (TERN). Original source: Charles Darwin University (NT) | 1.00 | 6 |
| field inventory | Joint Remote Sensing Research Program (2016): Biomass Plot Library - National collation of stem inventory data and biomass estimation, Australian field sites. Version 1.0.0. Terrestrial Ecosystem Research Network (TERN). Original source: University of Queensland | 0.04 | 70 |
| field inventory | Joint Remote Sensing Research Program (2016): Biomass Plot Library - National collation of stem inventory data and biomass estimation, Australian field sites. Version 1.0.0. Terrestrial Ecosystem Research Network (TERN). Original source: OEH NSW | 0.10 | 27 |

| | | | |
|---|---|---|---|
| field inventory | Joint Remote Sensing Research Program (2016): Biomass Plot Library - National collation of stem inventory data and biomass estimation, Australian field sites. Version 1.0.0. Terrestrial Ecosystem Research Network (TERN). Original source: University of NSW | 0.08 | 16 |
| field inventory | Condit et al. 2012. Barro Colorado Forest Census Plot Data, 2012 Version. DOI http://dx.doi.org/10.5479/data.bci.20130603. | 50.00 | 1 |
| field inventory | Gilbert & Jones 2024. Stem data from first three censuses on the University of California Santa Cruz Forest Ecology Research Plot [Dataset]. Dryad. https://doi.org/10.5061/dryad.6q573n64s | 16.00 | 1 |
| field inventory | Gonzalez-Akre et al. 2020. SCBI-ForestGEO/SCBI-ForestGEO-Data: first release with hydraulic traits data (v1.3). Zenodo. https://doi.org/10.5281/zenodo.4070038 | 25.00 | 1 |
| field inventory | Peijian et al. 2023. Forest survey data of a landscape in Pine Mountain [Dataset]. Dryad. https://doi.org/10.5061/dryad.h9w0vt4np | 40.00 | 1 |
| field inventory | Davis 2021. Twenty-five years of tree demography in a frequently burned oak woodland [Dataset]. Dryad. https://doi.org/10.5061/dryad.69p8cz92g | 16.00 | 1 |
| field inventory | Lakkana et al. (2022), Tropical montane forest in South Asia: Composition, structure and dieback in relation to soils and topography. Dryad Dataset. https://doi.org/10.5061/dryad.70rxwdbxv | 0.90 | 1 |
| field inventory | Pelissier et al. 2012. Tree demography in an undisturbed Dipterocarp permanent sample plot at Uppangala, Western Ghats of India. Ecology, 92: 1376-1376. doi:10.1890/10-1991.1 | 5.07 | 1 |
| field inventory | Kindermann et al. 2022: Dataset on Woody Aboveground Biomass, Disturbance Losses, and Wood Density from an African Savanna Ecosystem, V5. Mendeley Data. doi: 10.17632/3cs85wd3gb.5 | 0.10 | 41 |
| field inventory | Flake et al. 2022: Not all trees can make a forest: tree species composition and competition control forest encroachment in a tropical savanna. Dryad Dataset. https://doi.org/10.5061/dryad.vq83bk3tm | 0.70 | 1 |
| field inventory | Packee et al. 2016. Cooperative Alaska Forest Inventory ver 16. Environmental Data Initiative. https://doi.org/10.6073/pasta/7906aa46e03632f7bebaae23a8b46c8e (Accessed 2024-04-20) | 0.12 | 233 |
| field inventory | Rich and Fournier 1999. BOREAS TE-23 Map Plot Data. Data set. Available on-line [http://www.daac.ornl.gov] from Oak Ridge National Laboratory Distributed Active Archive Center, Oak Ridge, Tennessee, U.S.A. https://doi.org/10.3334/ORNLDAAC/359. | 0.30 | 6 |
| field inventory | Joint Remote Sensing Research Program (2016): Biomass Plot Library - National collation of stem inventory data and biomass estimation, Australian field sites. Version 1.0.0. Terrestrial Ecosystem Research Network (TERN). Original source: Flinders University (SA) | 0.25 | 15 |
| pre-aggregated | Ploton et al. 2020: A map of African humid tropical forest aboveground biomass derived from management inventories. Scientific Data 7, 221. | 1.50 | 55963 |
| pre-aggregated | Mitchard et al. 2014. Markedly divergent estimates of Amazon forest carbon density from ground plots and satellites. Global Ecology and Biogeography. doi: 10.1111/geb.12168. | 1.00 | 187 |

| | | | |
|---|---|---|---|
| pre-aggregated | Lewis et al. 2013. Above-ground biomass and structure of 260 African tropical forests. Philosophical Transactions of the Royal Society B: Biological Sciences 368 (1625):20120295. doi:10.1098/rstb.2012.0295. AfriTRON network data. | 1.00 | 218 |
| pre-aggregated | Colgan et al. 2013. Harvesting tree biomass at the stand level to assess the accuracy of field and airborne biomass estimation in savannas. Ecological Applications 23, 1170–1184. http://www.jstor.org/stable/23441615. Reported in Chave et al. 2014. | 2.12 | 1 |
| pre-aggregated | Schepaschenko et al. 2019: A global reference dataset for remote sensing of forest biomass. The Forest Observation System approach. DOI: 10.22022/ESM/03-2019.38 | 0.25 | 65 |
| pre-aggregated | Cuni-Sanchez A, White LJT, Calders K, Jeffery KJ, Abernethy K, Burt A, et al. (2016) African Savanna-Forest Boundary Dynamics: A 20-Year Study. PLoS ONE 11(6): e0156934. https://doi.org/10.1371/journal.pone.0156934 | 0.08 | 16 |
| pre-aggregated | Schepaschenko et al. 2017: A dataset of forest biomass structure for Eurasia. Scientific Data 4, 170070. | | 531 |
| pre-aggregated | Qie et al. 2017. Long-term carbon sink in Borneo’s forests halted by drought and vulnerable to edge effects. Nat Commun 8, 1966. https://doi.org/10.1038/s41467-017-01997-0. | 0.75 | 71 |
| pre-aggregated | Duque et al. 2021. Andean forests as globally important carbon sinks and future carbon refuges. Nat Commun 12, 3617 (2021). https://doi.org/10.1038/s41467-021-23955-7 | 1.00 | 119 |
| pre-aggregated | Williams et al. 2008. Carbon sequestration and biodiversity of re-growing miombo woodlands in Mozambique. Forest Ecology and Management 254, p. 145-155. | 3.50 | 1 |
| pre-aggregated | Sungpalee et al. 2009. Intra- and interspecific variation in wood density and fine-scale spatial distribution of stand-level wood density in a northern Thai tropical montane forest. Journal of Tropical Ecology 2009, 359-370. | 15.00 | 1 |

**Table S8. Sources of plot data for wood density mapping.** Shown are the sources that we used to derive community-weighted wood density estimates across the globe, their median area (in ha), and the total number of plots. We divide plot data into four broad categories: national forest inventories and similar large-scale plot networks (NFI), local field inventories at specific study sites, phytosociological studies that typically only provide (relative) basal area by species, and pre-aggregated data that directly provide values for community-weighted wood density.

| *species* | *family* | *lineage* | *samples* | *WD avg (g cm⁻³)* | *WD lmer (g cm⁻³)* |
|---|---|---|---|---|---|
| Adansonia digitata | Malvaceae | Eudicots | 2 | 0.32 | 0.31 |
| Adansonia fony var. rubrostipa | Malvaceae | Eudicots | 2 | 0.29 | 0.25 |
| Adansonia grandidieri | Malvaceae | Eudicots | 1 | 0.09 | 0.17 |
| Adansonia madagascariensis | Malvaceae | Eudicots | 2 | 0.19 | 0.21 |
| Adansonia za | Malvaceae | Eudicots | 2 | 0.3 | 0.26 |
| Adenium obesum | Apocynaceae | Eudicots | 1 | 0.17 | 0.26 |
| Alluaudia ascendens | Didiereaceae | Eudicots | 2 | 0.38 | 0.38 |
| Alluaudia comosa | Didiereaceae | Eudicots | 1 | 0.4 | 0.38 |
| Alluaudia dumosa | Didiereaceae | Eudicots | 10 | 0.42 | 0.4 |
| Alluaudia humbertii | Didiereaceae | Eudicots | 1 | 0.46 | 0.41 |
| Alluaudia procera | Didiereaceae | Eudicots | 29 | 0.28 | 0.27 |
| Alluaudiopsis fiherenensis | Didiereaceae | Eudicots | 1 | 0.51 | 0.49 |
| Anredera cordifolia | Basellaceae | Eudicots | 2 | 0.2 | 0.27 |
| Asclepias subulata | Apocynaceae | Eudicots | 1 | 0.31 | 0.37 |
| Brachychiton australis | Malvaceae | Eudicots | 2 | 0.38 | 0.39 |
| Bursera fagaroides | Burseraceae | Eudicots | 4 | 0.36 | 0.36 |
| Bursera filicifolia | Burseraceae | Eudicots | 1 | 0.4 | 0.4 |
| Bursera microphylla | Burseraceae | Eudicots | 1 | 0.3 | 0.35 |
| Carnegiea gigantea | Cactaceae | Eudicots | 1 | 0.3 | 0.34 |
| Cavanillesia platanifolia | Malvaceae | Eudicots | 8 | 0.2 | 0.22 |
| Ceiba insignis | Malvaceae | Eudicots | 10 | 0.27 | 0.3 |
| Cereus jamacaru | Cactaceae | Eudicots | 1 | 0.3 | 0.34 |

| | | | | | |
|---|---|---|---|---|---|
| Cissus aralioides | Vitaceae | Eudicots | 1 | 0.45 | 0.41 |
| Cissus repens | Vitaceae | Eudicots | 1 | 0.36 | 0.35 |
| Cylindropuntia acanthocarpa | Cactaceae | Eudicots | 1 | 0.67 | 0.64 |
| Cylindropuntia cholla | Cactaceae | Eudicots | 1 | 0.52 | 0.55 |
| Decarya madagascariensis | Didiereaceae | Eudicots | 1 | 0.55 | 0.52 |
| Delonix boiviniana | Fabaceae | Eudicots | 1 | 0.34 | 0.37 |
| Delonix leucantha | Fabaceae | Eudicots | 2 | 0.3 | 0.33 |
| Didierea madagascariensis | Didiereaceae | Eudicots | 2 | 0.5 | 0.48 |
| Didierea trollii | Didiereaceae | Eudicots | 1 | 0.43 | 0.44 |
| Entandrophragma caudatum | Meliaceae | Eudicots | 4 | 0.62 | 0.55 |
| Erythrina flabelliformis | Fabaceae | Eudicots | 1 | 0.13 | 0.22 |
| Euphorbia abyssinica | Euphorbiaceae | Eudicots | 1 | 0.31 | 0.36 |
| Euphorbia californica | Euphorbiaceae | Eudicots | 1 | 0.36 | 0.38 |
| Euphorbia candelabrum | Euphorbiaceae | Eudicots | 3 | 0.26 | 0.3 |
| Euphorbia grandifolia | Euphorbiaceae | Eudicots | 2 | 0.26 | 0.31 |
| Euphorbia neriifolia | Euphorbiaceae | Eudicots | 1 | 0.34 | 0.36 |
| Euphorbia stenoclada | Euphorbiaceae | Eudicots | 12 | 0.39 | 0.38 |
| Euphorbia tirucalli | Euphorbiaceae | Eudicots | 28 | 0.4 | 0.39 |
| Fouquieria diguetii | Fouquieriaceae | Eudicots | 1 | 0.33 | 0.39 |
| Fouquieria formosa | Fouquieriaceae | Eudicots | 1 | 0.38 | 0.4 |
| Fouquieria splendens | Fouquieriaceae | Eudicots | 1 | 0.53 | 0.49 |
| Hoya australis | Apocynaceae | Eudicots | 5 | 0.55 | 0.58 |
| Jatropha cinerea | Euphorbiaceae | Eudicots | 1 | 0.2 | 0.25 |
| Jatropha cuneata | Euphorbiaceae | Eudicots | 1 | 0.32 | 0.31 |

| | | | | | |
|---|---|---|---|---|---|
| Jatropha curcas | Euphorbiaceae | Eudicots | 2 | 0.23 | 0.24 |
| Jatropha gaumeri | Euphorbiaceae | Eudicots | 1 | 0.43 | 0.31 |
| Jatropha mahafalensis | Euphorbiaceae | Eudicots | 1 | 0.14 | 0.2 |
| Jatropha malacophylla | Euphorbiaceae | Eudicots | 2 | 0.28 | 0.25 |
| Jatropha peltata | Euphorbiaceae | Eudicots | 1 | 0.41 | 0.34 |
| Leptocereus paniculatus | Cactaceae | Eudicots | 1 | 0.5 | 0.49 |
| Leucostele terscheckii | Cactaceae | Eudicots | 1 | 0.25 | 0.32 |
| Leuenbergeria guamacho | Cactaceae | Eudicots | 2 | 0.54 | 0.51 |
| Leuenbergeria marcanoi | Cactaceae | Eudicots | 1 | 0.7 | 0.58 |
| Leuenbergeria portulacifolia | Cactaceae | Eudicots | 1 | 0.6 | 0.54 |
| Leuenbergeria quisqueyana | Cactaceae | Eudicots | 1 | 0.62 | 0.55 |
| Neoraimondia herzogiana | Cactaceae | Eudicots | 1 | 0.52 | 0.52 |
| Operculicarya decaryi | Anacardiaceae | Eudicots | 1 | 0.2 | 0.29 |
| Opuntia excelsa | Cactaceae | Eudicots | 1 | 0.3 | 0.37 |
| Pachycereus pecten-aboriginum | Cactaceae | Eudicots | 1 | 0.4 | 0.35 |
| Pachycereus pringlei | Cactaceae | Eudicots | 1 | 0.25 | 0.3 |
| Pachypodium lamerei | Apocynaceae | Eudicots | 11 | 0.1 | 0.1 |
| Pereskia diaz-romeroana | Cactaceae | Eudicots | 1 | 0.61 | 0.53 |
| Pereskia sacharosa | Cactaceae | Eudicots | 1 | 0.57 | 0.51 |
| Pereskia weberiana | Cactaceae | Eudicots | 1 | 0.64 | 0.55 |
| Phytolacca dioica | Phytolaccaceae | Eudicots | 2 | 0.23 | 0.3 |
| Pilosocereus pachycladus | Cactaceae | Eudicots | 1 | 0.53 | 0.48 |
| Pilosocereus purpusii | Cactaceae | Eudicots | 1 | 0.39 | 0.43 |
| Pittocaulon praecox | Asteraceae | Eudicots | 2 | 0.44 | 0.46 |

| | | | | | |
|---|---|---|---|---|---|
| **Pseudobombax ellipticum** | Malvaceae | Eudicots | 6 | 0.35 | 0.33 |
| **Pseudobombax grandiflorum** | Malvaceae | Eudicots | 6 | 0.33 | 0.34 |
| **Sedum oxypetalum** | Crassulaceae | Eudicots | 2 | 0.1 | 0.15 |
| **Steganotaenia araliacea** | Apiaceae | Eudicots | 2 | 0.46 | 0.48 |
| **Stenocereus chrysocarpus** | Cactaceae | Eudicots | 1 | 0.39 | 0.45 |
| **Stenocereus thurberi** | Cactaceae | Eudicots | 1 | 0.49 | 0.49 |
| **Sterculia africana** | Malvaceae | Eudicots | 4 | 0.49 | 0.46 |
| **Sterculia murex** | Malvaceae | Eudicots | 2 | 0.54 | 0.48 |
| **Talinum paniculatum** | Talinaceae | Eudicots | 1 | 0.68 | 0.69 |
| **Cordyline australis** | Asparagaceae | Monocots | 2 | 0.28 | 0.34 |
| **Yucca baccata** | Asparagaceae | Monocots | 1 | 0.27 | 0.35 |
| **Yucca brevifolia** | Asparagaceae | Monocots | 2 | 0.42 | 0.43 |
| **Yucca elata** | Asparagaceae | Monocots | 1 | 0.39 | 0.41 |
| **Yucca treculeana** | Asparagaceae | Monocots | 1 | 0.58 | 0.51 |

**Table S9. List of succulents in the GWDD v.2.** Show are all species in the GWDD v.2 that are classified as succulents. Shown are their family, lineage (eudicots vs. monocots), the number of samples in the GWDD v.2, the average value of raw measurements, and a refined estimate of species means via linear mixed effects modelling that considers taxonomic structure (Fischer et al., 2025.). Classification as succulents is based on records from the Sukkulentensammlung Zürich, Switzerland (Anderson & Eggli, 2005; Eggli, 2004; Eggli & Nyffeler, 2020, 2023)

#### B. Supplementary Figures

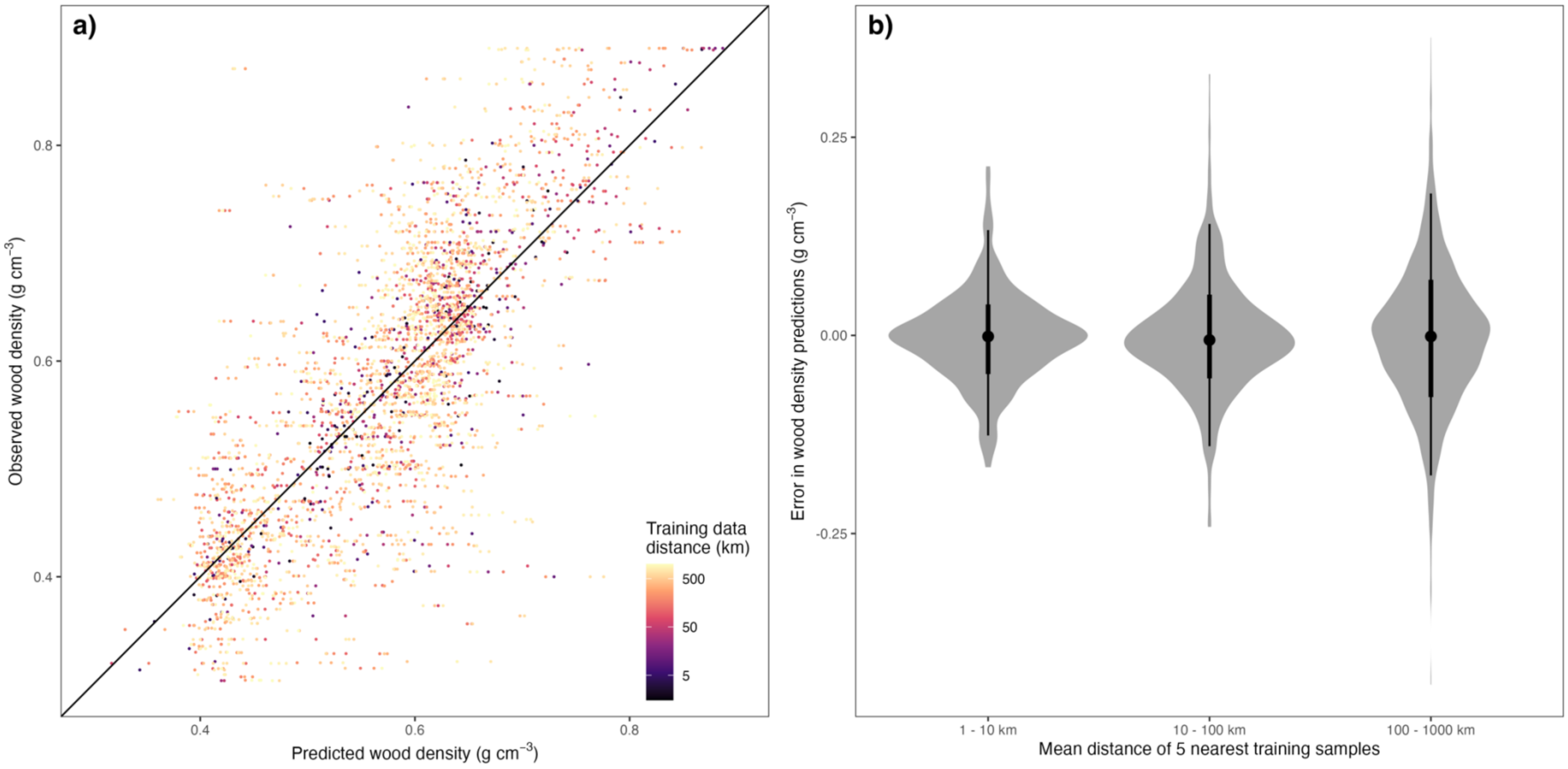

**Fig. S1. Spatial leave-one-out cross validation.** Shown are the results of a spatial leave-one-out cross validation, using the 1,000 validation plots shown in Fig. S1. We adapted the methodology described in Ploton et al., 2020 in the following way: for each of the 1,000 plots, we predicted community-weighted wood density four times: once by leaving out the plot itself and predicting its wood density with a random forest model trained on the remaining 999 plots, and an additional three time by choosing a random distance between 0 and 1000 km and, removing all training data within this “exclusion radius” before fitting the random forest model. We removed duplicates, where the exclusion radius is smaller than the validation plot’s original distance to the nearest training data and data points with distances > 1000 km, leaving a total of 3,841 data points for validation. Panel a) shows predicted wood densities and observed wood densities, coloured by the distance of validation plots to the five nearest training samples, panel b) shows the distribution of prediction errors as function of the distance to training samples, loosely grouped into 3 logarithmic bins (1 – 10 km, 10 – 100 km and 100 – 1000 km).

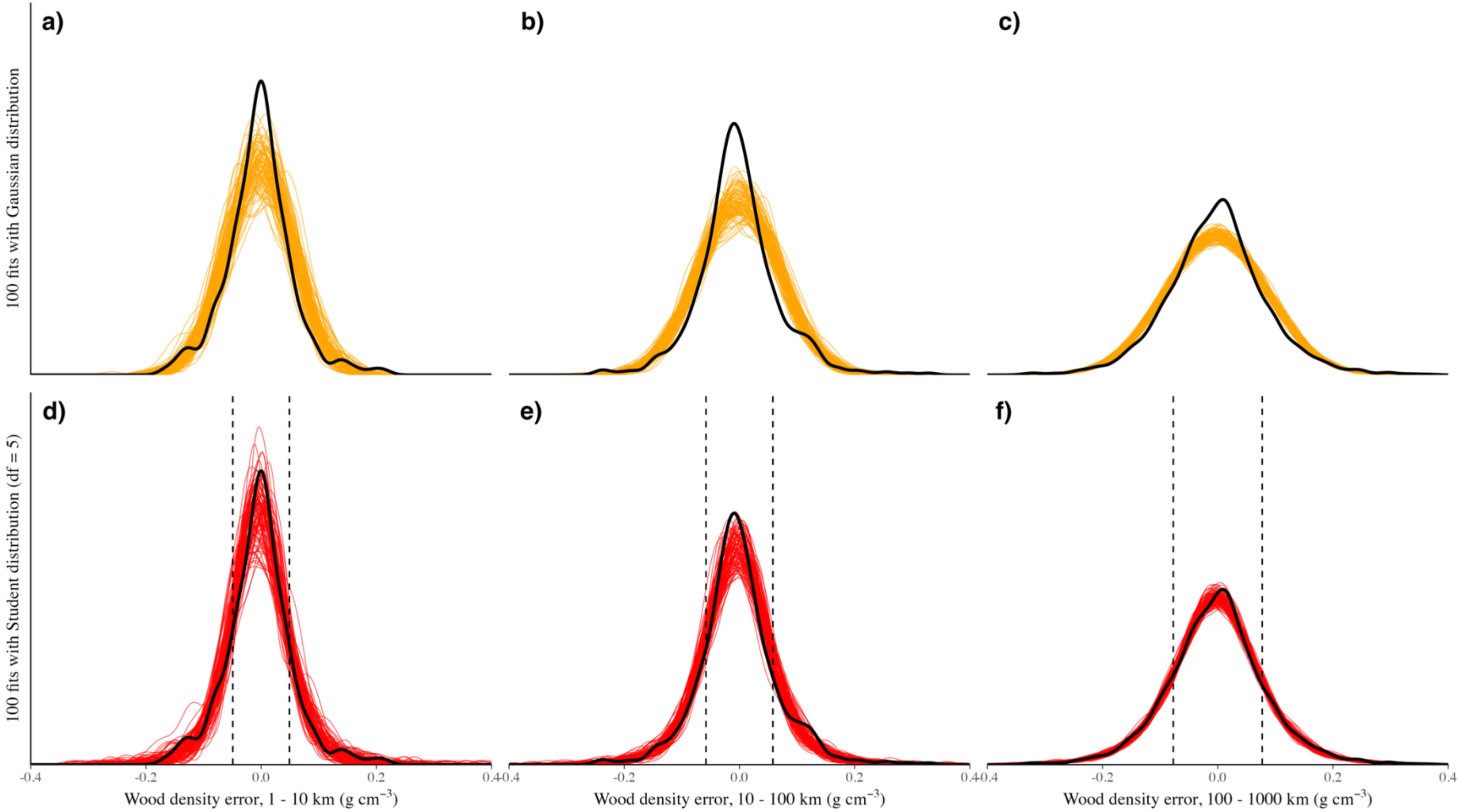

**Fig. S2. Best-fit error distribution ("posterior predictive check").** Shown are comparisons between the observed distribution of wood density errors (solid black lines), and either 100 fits of Gaussian distributions (panels a-c) or 100 fits of location-scale student distributions with 5 degrees of freedom (panels d-f). As in Fig. S1b, validation data are loosely grouped according to distance from nearest training data: 1-10 km (panels a and d), 10-100 km (panels b and e) and 100-1000 km (panels c and f). The dashed vertical lines in panels d-f show the $\sigma_t$* metric where $\sigma_t$* = 1.11 x $\sigma_t$ and $\sigma_t$ is the variance parameter of the student distribution. If data points follow a student distribution with 5 degrees of freedom, then ~68.3% of data points will be within the interval [-$\sigma_t$*, $\sigma_t$*]. The $\sigma_t$* metric can thus be seen as roughly equivalent of the standard deviation of a Gaussian distribution, which also covers ~68.3% of data points in its lower and upper bound. Gaussian and Student distributions were fit with the brms package, and the posterior draws were plotted via the inbuilt pp_check() function. Note that the grouping into distance classes is just for visualization purposes – in the actual models, error distributions were fit directly as a function of the (continuous) distance to the nearest training data.

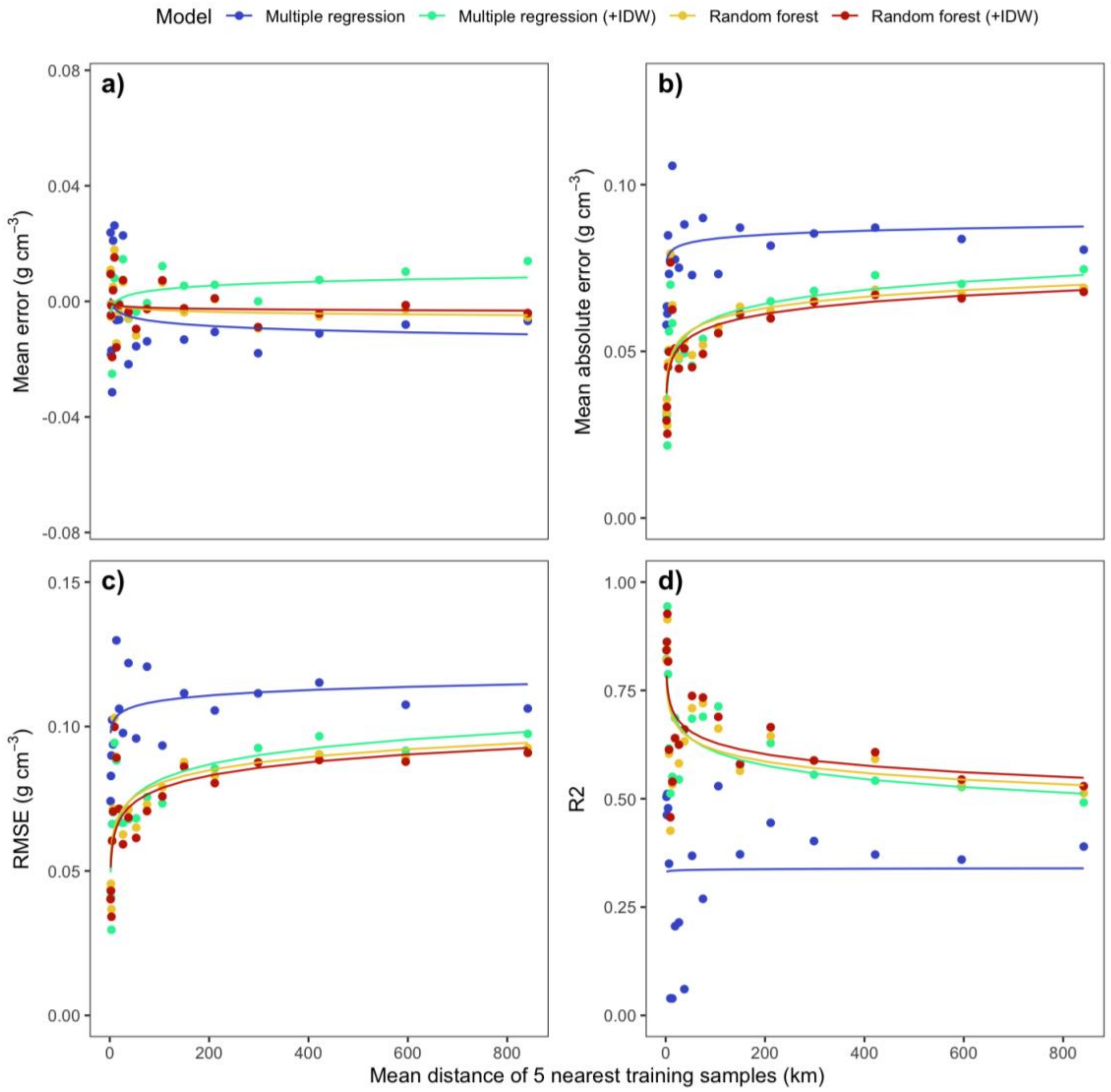

**Fig. S3. Predictive power of models with distance from training data (original scales).** Shown are the predictive performances of four models for wood density mapping, in relation to the distance from the five nearest training samples. The approaches comprise a multiple regression and a random forest model, in each case with or without inverse-distance weighted (IDW) interpolation of residuals and separated by colours. The four evaluation metrics (panels a-d) are calculated based on spatial leave-one-out cross validation on 1,000 plots (Fig. S1), whereby each plot's wood density is predicted four times: once by leaving out the plot and training the models on the remaining 999 plots, and an additional three time by choosing a random distance between 0 and 1000 km and, removing all training data within this "exclusion radius" before training the models. We removed duplicates, where the exclusion radius was smaller than the validation plot's original distance to the nearest training samples, as well as data points with distances > 1000 km, leaving a total of 3,841 data points for validation. For plotting, points were sorted into 20 distance bins on log-scale, and each metric's relationship to the distance from the nearest training samples was modelled as a linear regression against the log-transformed distance. Note how in all models, except the simplest (multiple) regression model, predictive power decreases rapidly with increasing distance to training samples up to a distance of ca. 100-200 km, and continuous to decrease even beyond that scale, albeit more slowly. This suggests that all three models are highly susceptible to spatial autocorrelation effects. Note that our approach for mapping wood density errors globally (Fig. 1b in main text, Fig. S5 below) is similar to the linear regressions employed here, but avoids binning, relies on a student response distribution and includes predicted wood density as covariate.

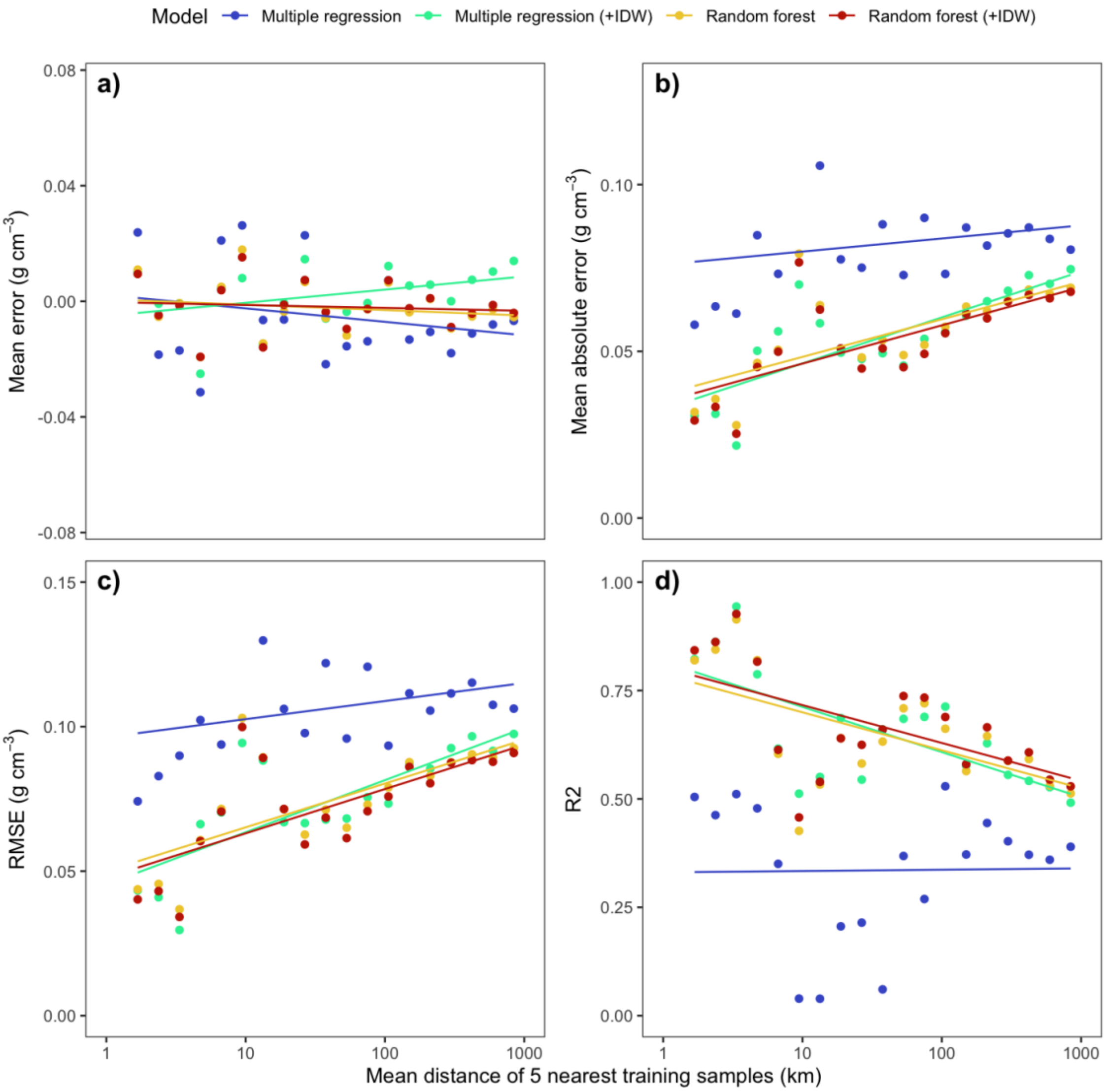

**Fig. S4. Predictive power of models with distance from training samples (log-scales).** Shown are the predictive performances of four modelling approaches for wood density mapping, in relation to distance from the nearest training samples. Same as Fig. S3, but with log-transformed axes.

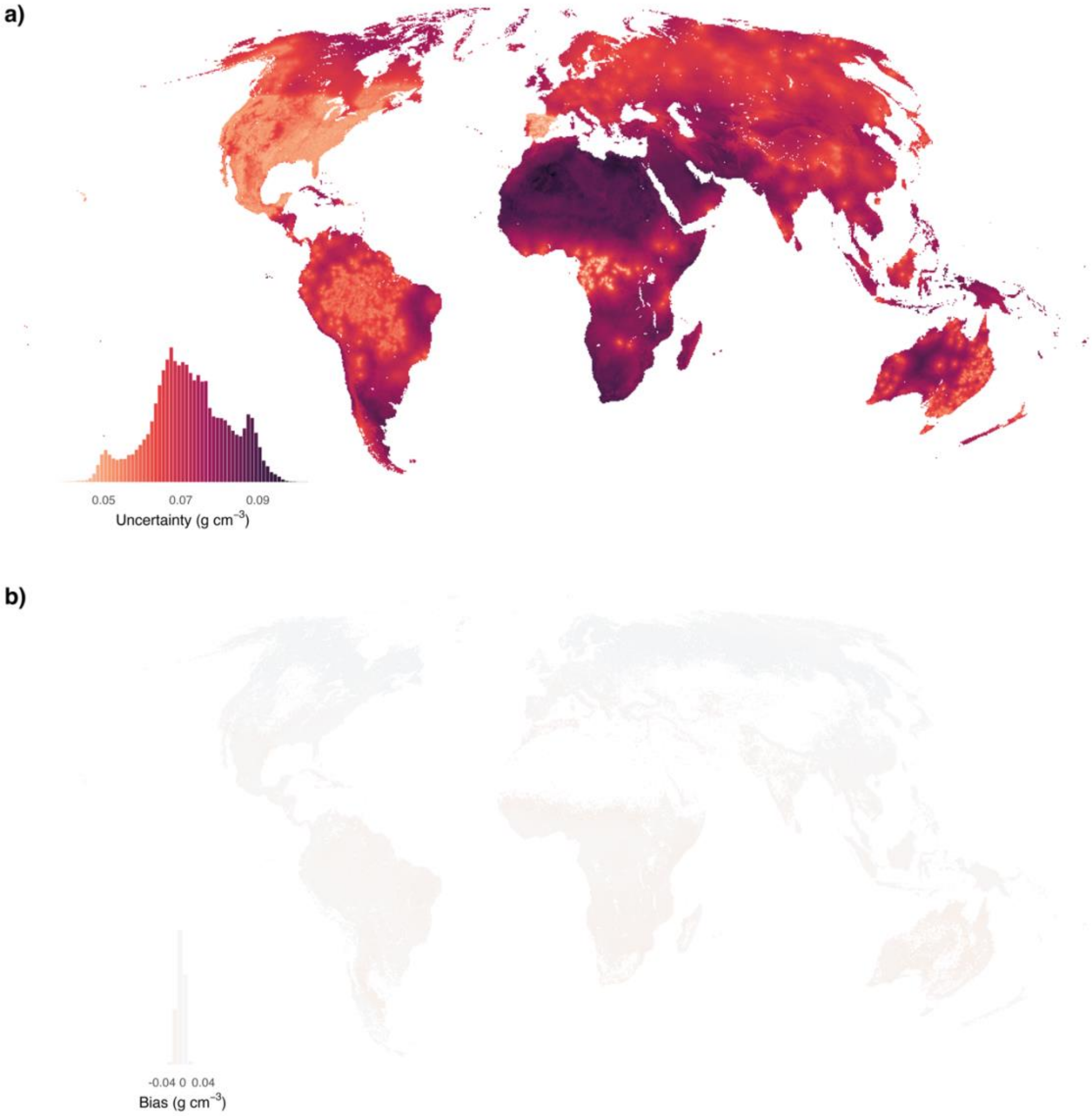

**Fig. S5. Global map of uncertainty and bias in wood density predictions.** The two maps are the equivalent of panels b) and c) in Fig. 1 in the main text, but with uncertainty (a) and bias (b) given in the original units instead of normalizing by the predicted wood density. The resolution is also 1 km resolution in equal-area Mollweide projection (ESRI:54009). The insets show both the colour scale and the histogram of values. The colourscale in panel b has been chosen so as to be comparable to the panel b colourscales in Figs. S7-10. Areas without woody vegetation (i.e., no significant tree or shrub cover) are masked out.

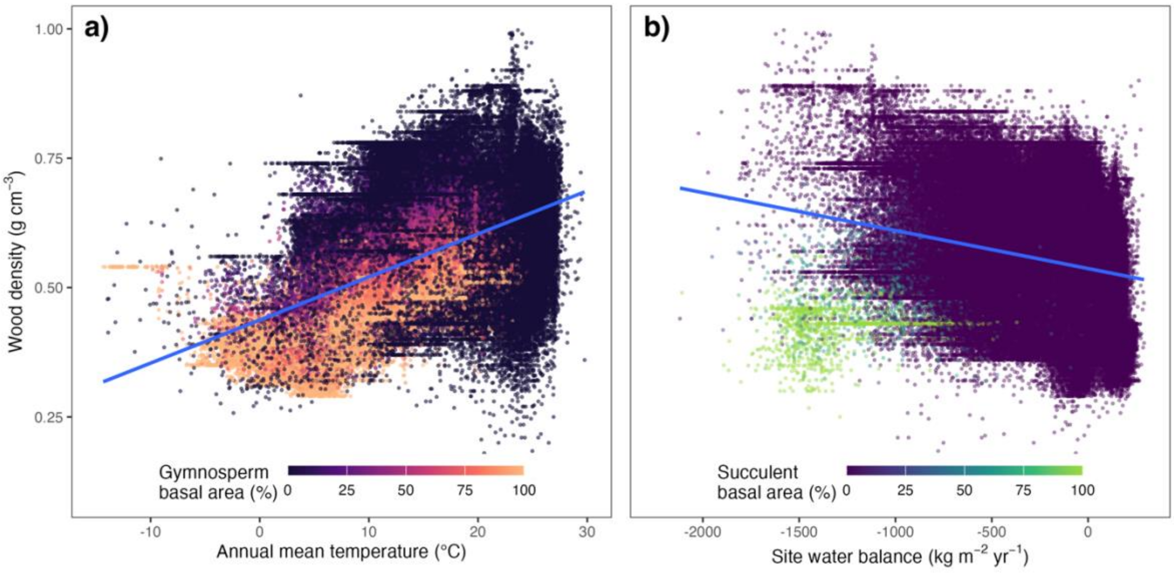

**Fig. S6 Environmental and biogeographic determinants of wood density (all plots).** This plots the same relationships as in Fig. 2 in the main text, but across all plots (n = 300,949), with points coloured by the proportion of gymnosperm basal area (panel a) or proportion of succulent basal area (panel b) in each plot. Note how succulent ecosystems are clear outliers in the negative relationship between site water balance and wood density.

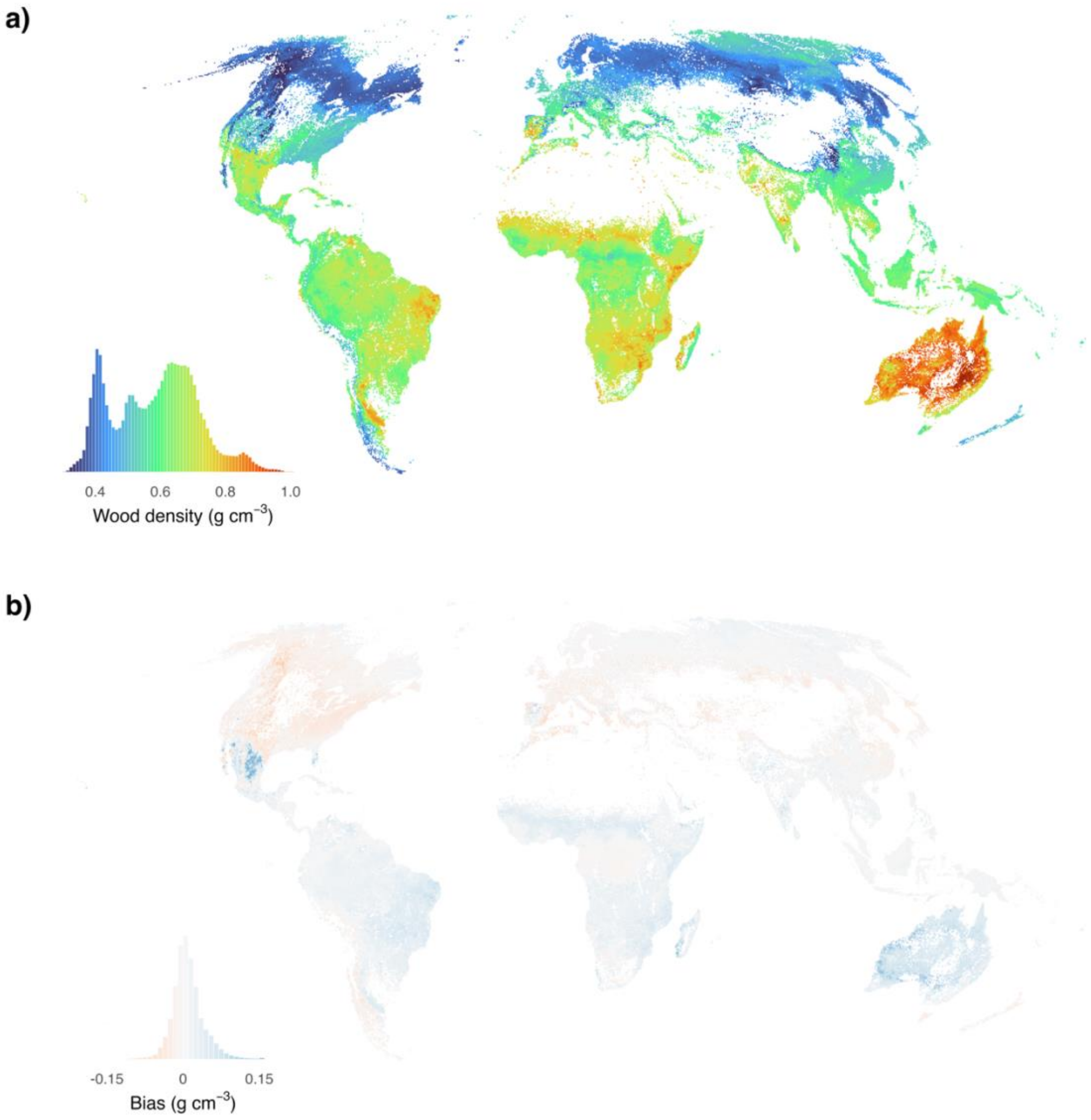

**Fig. S7. Comparison with map based on original version of GWDD.** Panel a shows the same map and histogram as in Fig. 1 in the main text, but computed with wood density records from the previous version of the Global Wood Density Database (GWDD). Panel b shows the bias with respect to the updated database, again both as map and as histogram. Bias was estimated as the difference from the reference map (Fig. 1a) minus the reference map's bias (Fig. S5b and Fig. 1c). The colourscale in panel b has been chosen so as to be comparable to the panel b colourscales in Figs. S5 and S8-10. Note that overall patterns in wood density are very similar, but that biases can be substantial regionally (up to or exceeding 0.1 g cm$^{-3}$).

a)

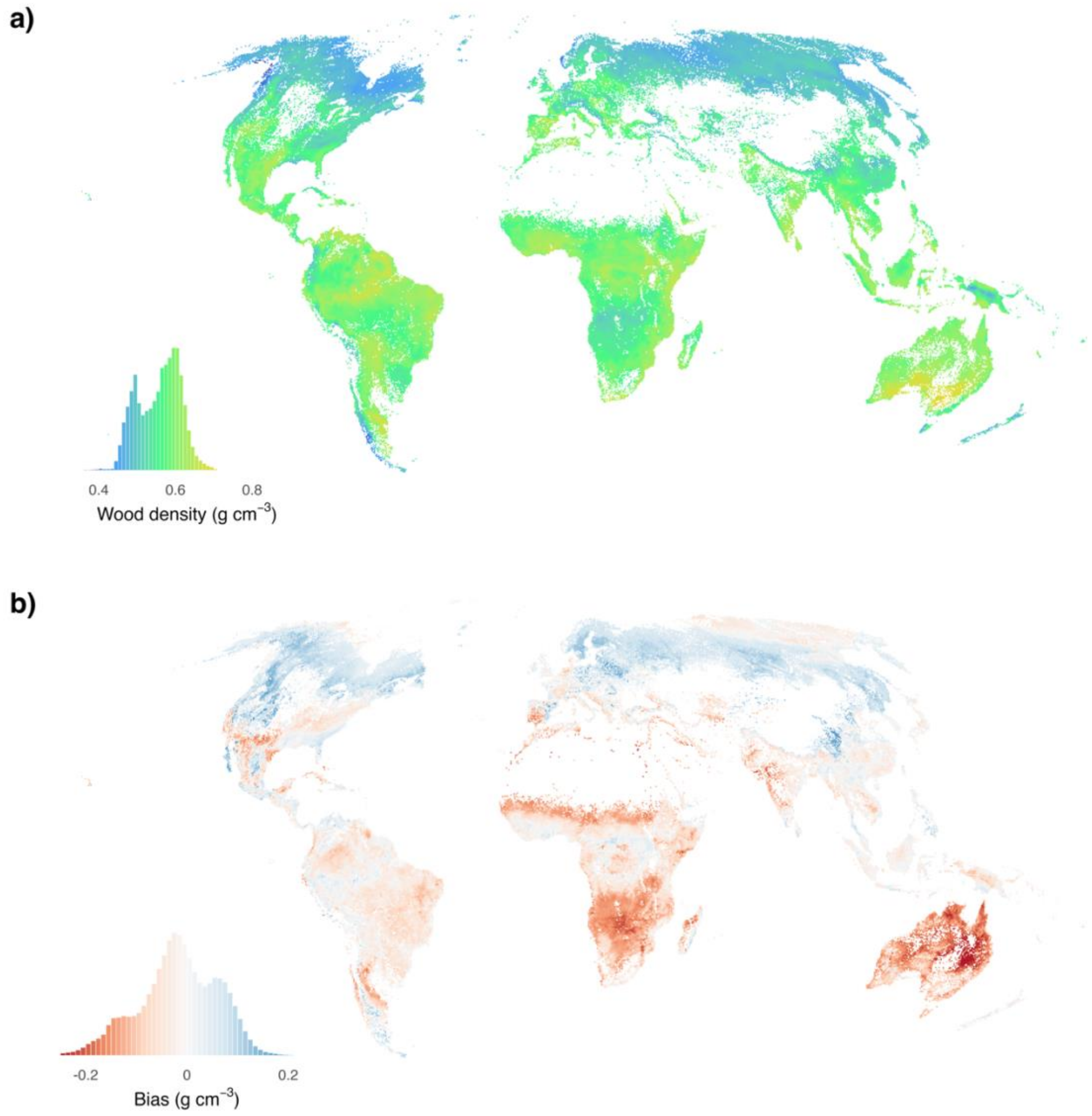

**Fig. S8. Comparison with map based on Boonman et al. 2020.** Panel a shows the same map and histogram as in Fig. 1a in the main text, but based on a reprojected (Mollweide) and resampled (1 km resolution) map provided in Boonman et al. 2020. The colourscale corresponds to the colourscale in Fig. 1a. Panel b shows the bias with respect to our reference map, again both as map and as histogram. Bias was estimated as the difference from the reference map (Fig. 1a) minus the reference map's bias (Fig. S5b and Fig. 1c).

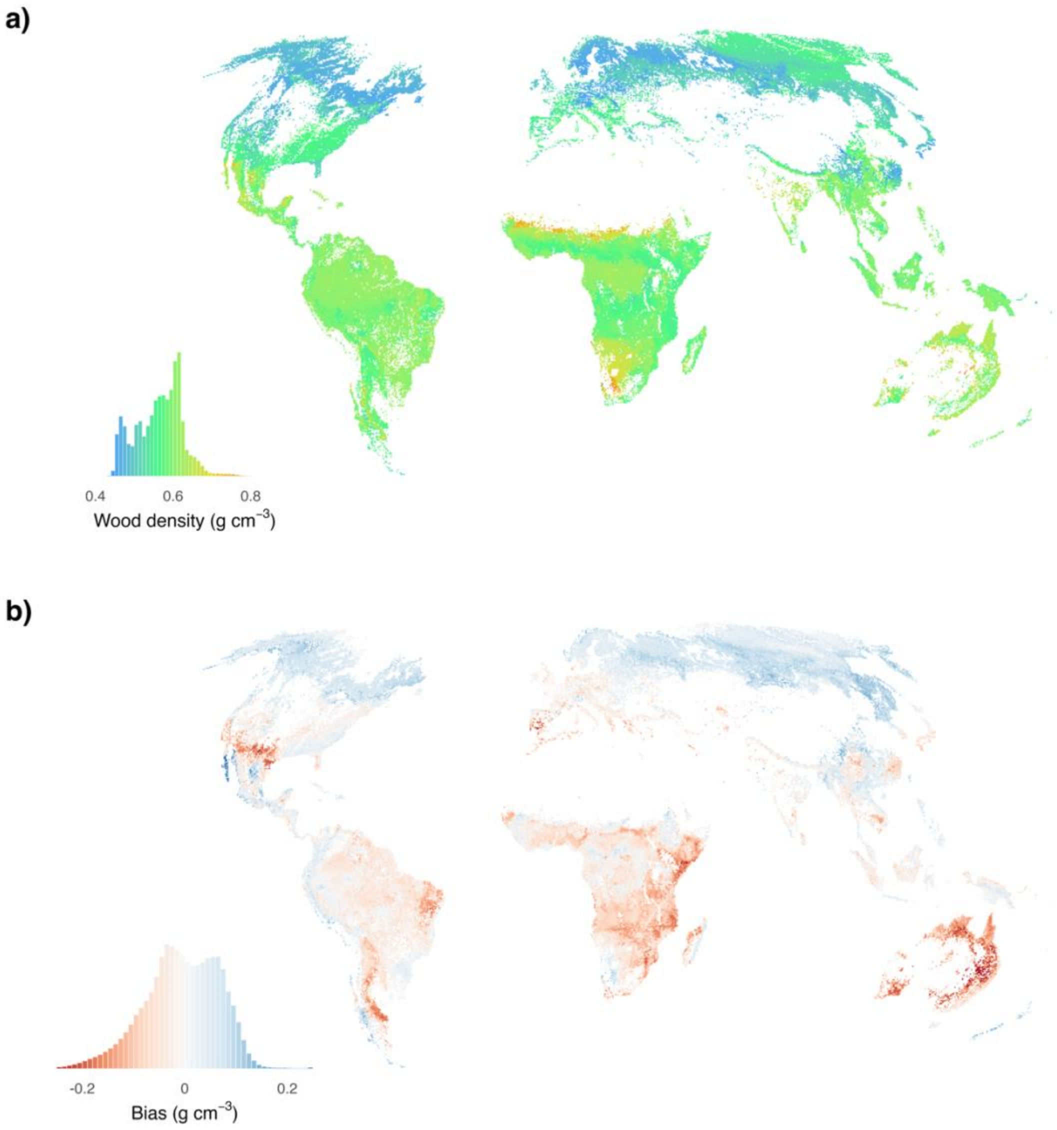

**Fig. S9. Comparison with map based on Yang et al. 2024.** Panel a shows the same map and histogram as in Fig. 1a in the main text, but based on a reprojected (Mollweide) and resampled (1 km resolution) map provided in Yang et al. 2024. The colourscale corresponds to the colourscale in Fig. 1a. Panel b shows the bias with respect to our reference map, again both as map and as histogram. Bias was estimated as the difference from the reference map (Fig. 1a) minus the reference map's bias (Fig. S5b and Fig. 1c).

a)

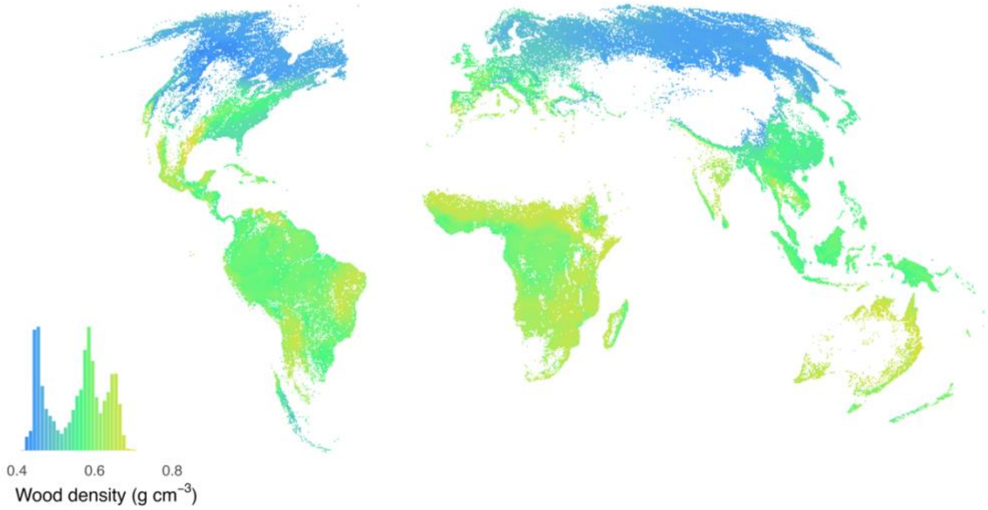

b)

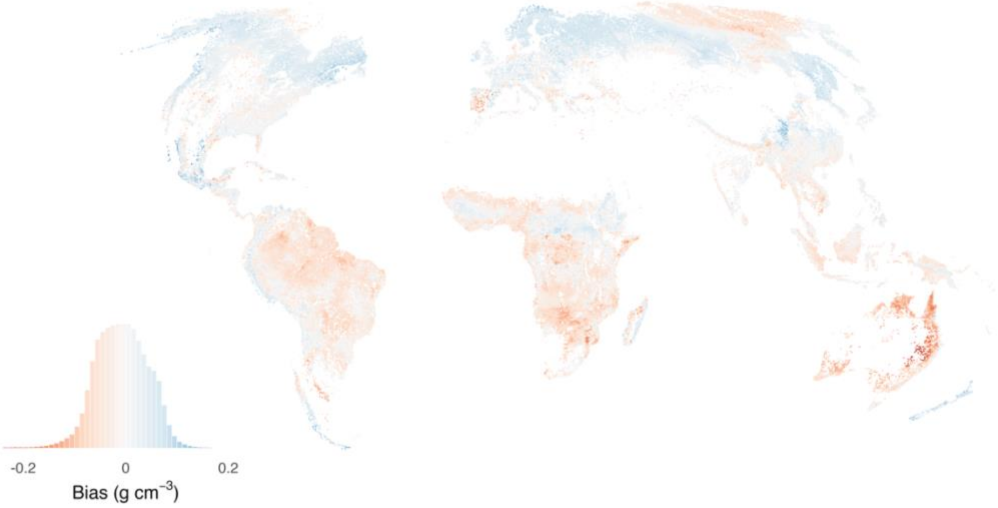

**Fig. S10. Comparison with map based on Mo et al. 2024.** Panel a shows the same map and histogram as in Fig. 1a in the main text, but based on a reprojected (Mollweide) and resampled (1 km resolution) map provided in Mo et al. 2024. The colourscale corresponds to the colourscale in Fig. 1a. Panel b shows the bias with respect to our reference map, again both as map and as histogram. Bias was estimated as the difference from the reference map (Fig. 1a) minus the reference map's bias (Fig. S5b and Fig. 1c).

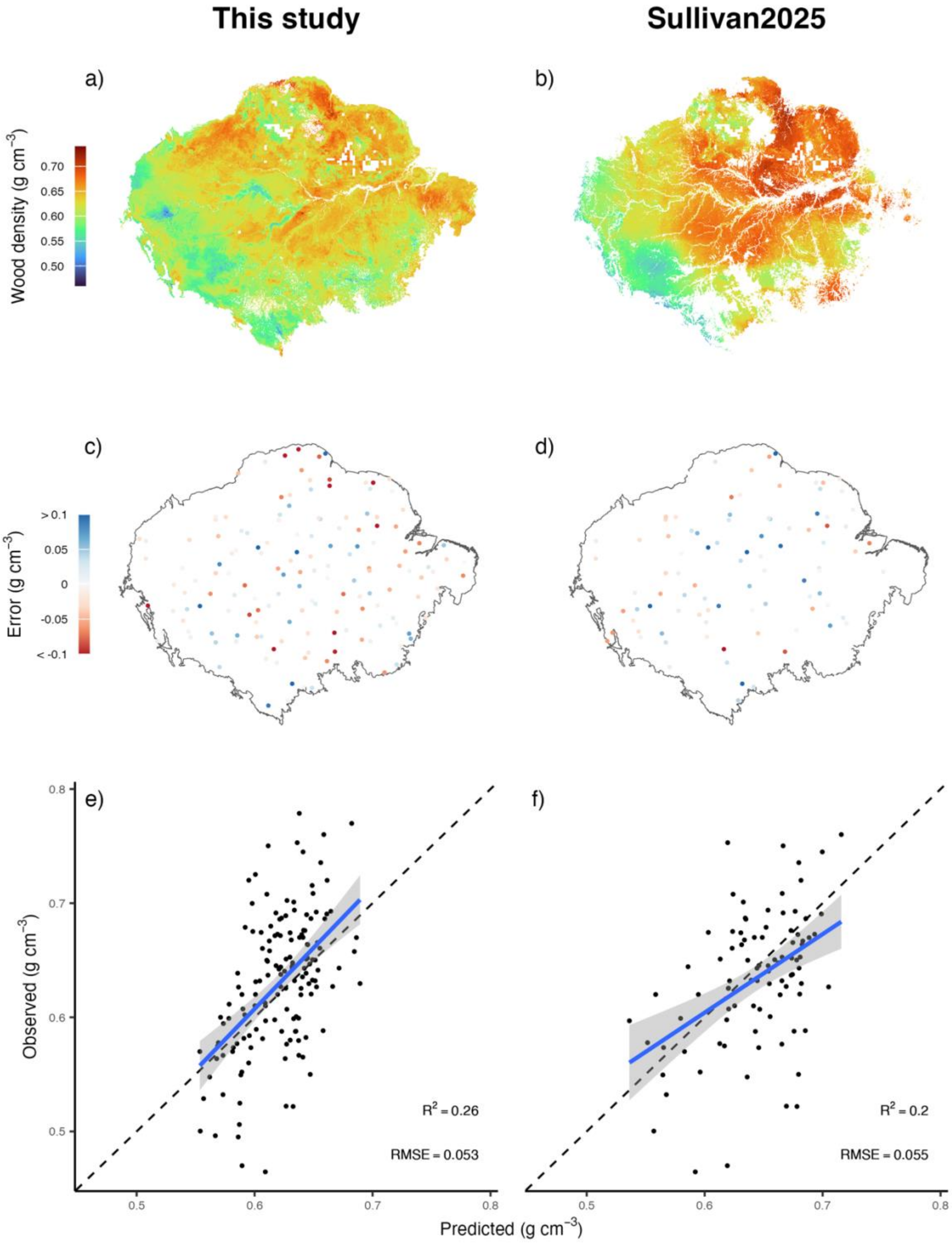

**Fig. S11. Wood density variation across Amazonia, comparing global and local maps.** Exactly the same panels as in Figure 3 in the main text, but comparing the global map produced in this study with a map produced for South American tropical forests (Sullivan et al., 2025).

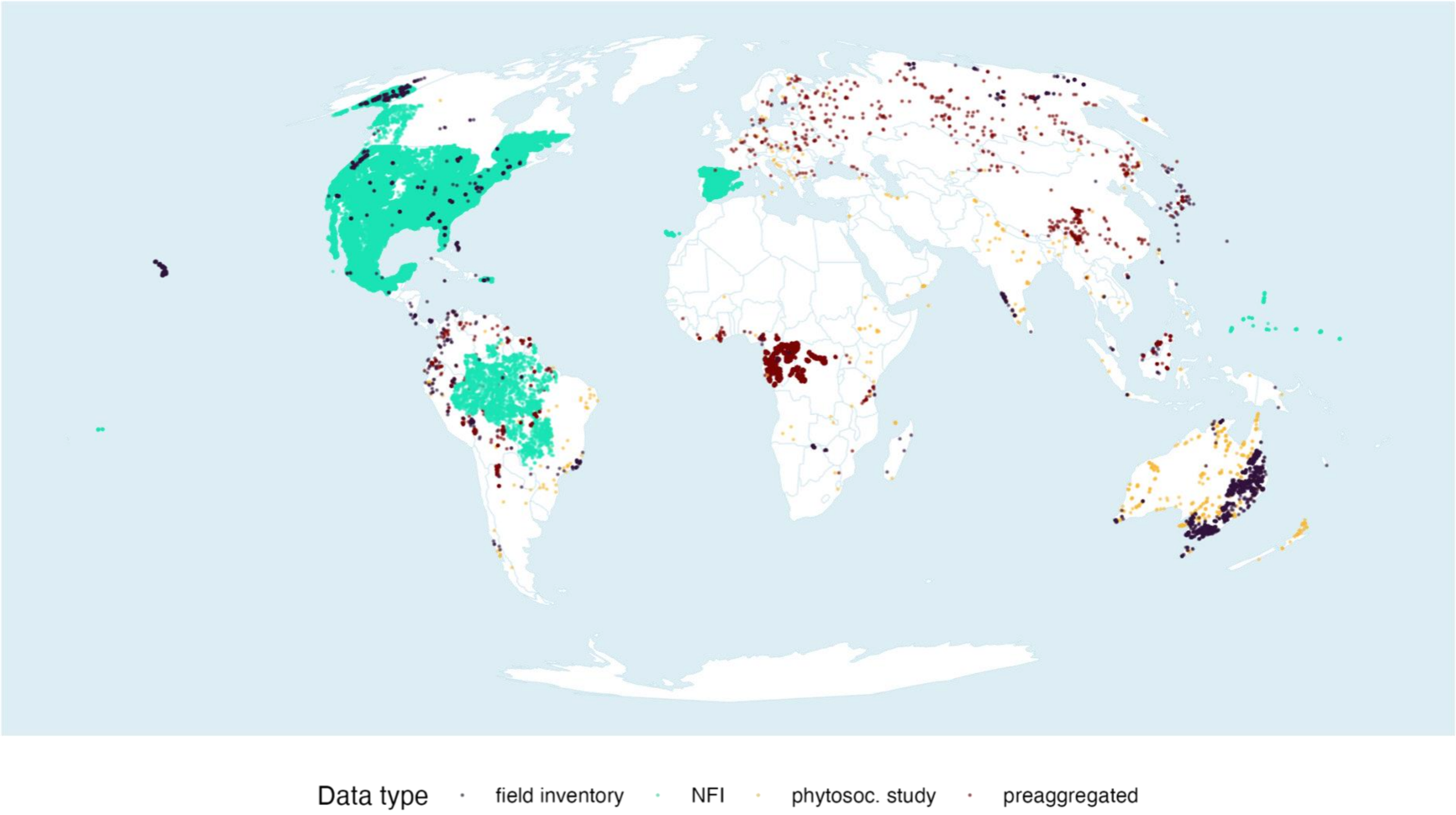

**Fig. S12. Location of plots used for wood density mapping.** Shown are the locations of all plots used as input for wood density mapping, separated by data type.

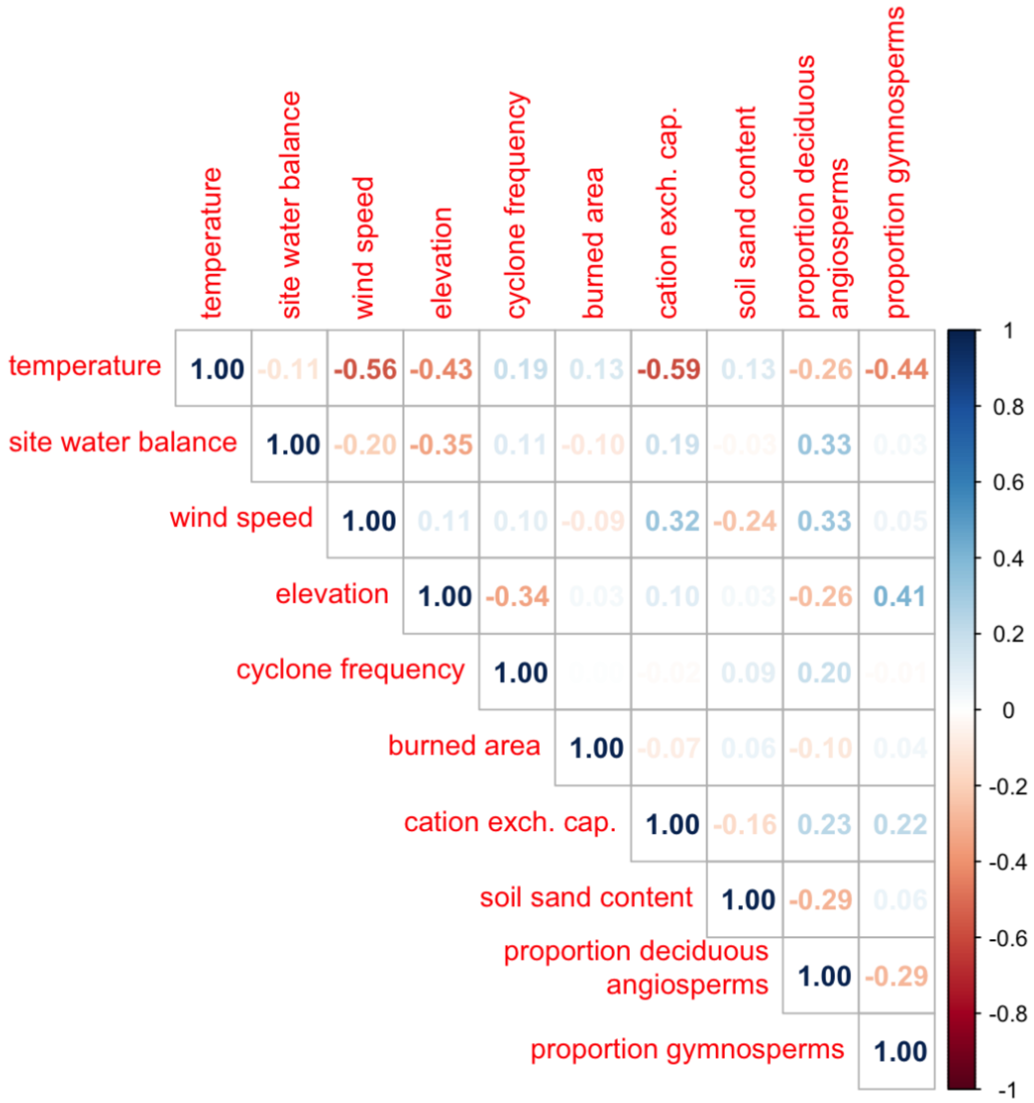

**Fig. S13. Correlation between predictors for wood density mapping.** Shown are the correlations between the 10 predictor layers. Note that values never exceed 0.6, which should minimize issues with covariance in modelling wood density.
